# Hyperosmotic stress induces downstream-of-gene transcription and alters the RNA Polymerase II interactome despite widespread transcriptional repression

**DOI:** 10.1101/2020.06.30.178103

**Authors:** Nicolle A. Rosa-Mercado, Joshua T. Zimmer, Maria Apostolidi, Jesse Rinehart, Matthew D. Simon, Joan A. Steitz

**Affiliations:** Department of Molecular Biophysics and Biochemistry, Yale University, New Haven, CT 06536, USA; Department of Cellular and Molecular Physiology, Yale University, New Haven, CT 06510, USA; Systems Biology Institute, Yale University, West Haven, CT 06477, USA; Howard Hughes Medical Institute, Yale University, New Haven, CT 06536, USA

## Abstract

Stress-induced readthrough transcription results in the synthesis of thousands of downstream-of-gene (DoG) containing transcripts. The mechanisms underlying DoG formation during cellular stress remain unknown. Nascent transcription profiles during DoG induction in human cell lines using TT-TimeLapse-seq revealed that hyperosmotic stress induces widespread transcriptional repression. Yet, DoGs are produced regardless of the transcriptional level of their upstream genes. ChIP-seq confirmed that the stress-induced redistribution of RNA Polymerase (Pol) II correlates with the transcriptional output of genes. Stress-induced alterations in the Pol II interactome are observed by mass spectrometry. While subunits of the cleavage and polyadenylation machinery remained Pol II-associated, Integrator complex subunits dissociated from Pol II under stress conditions. Depleting the catalytic subunit of the Integrator complex, Int11, using siRNAs induces hundreds of readthrough transcripts, whose parental genes partially overlap those of stress-induced DoGs. Our results provide insights into the mechanisms underlying DoG production and how Integrator activity influences DoG transcription.

**In brief:** Rosa-Mercado et al. report that hyperosmotic stress causes widespread transcriptional repression in human cells, yet DoGs arise regardless of the transcriptional response of their upstream genes. They find that the interaction between Pol II and Integrator is disrupted by hypertonicity and that knocking down the Integrator nuclease leads to DoG production.

**Highlights:**

- Hyperosmotic stress triggers transcriptional repression of many genes.
- DoG RNAs arise independent of the transcriptional level of their upstream gene.
- The interaction between Pol II and Integrator subunits decreases after salt stress.
- Depletion of the Int11 nuclease subunit induces the production of hundreds of DoGs.

## Introduction

Conditions of cellular stress, such as hyperosmotic stress, heat shock, oxidative stress, viral infections and renal cancer lead to readthrough transcription (Vilborg et al. 2015; Vilborg et al. 2017; Rutkowski et al. 2015; Bauer et al. 2018; Grosso et al. 2015; reviewed in Vilborg and Steitz 2017). This phenomenon is defined as transcription extending past the annotated termination sites of genes. This stress-induced transcription results in the production of thousands of long noncoding RNAs (lncRNAs), known as downstream-of-gene (DoG) containing transcripts. DoGs arise minutes after stress from approximately 10% of human protein-coding genes and are continuous with their upstream mRNAs. DoG RNAs are defined as extending over 5 kilobases (kb) past the normal termination site of their gene of origin. These transcripts can be over 200 kb long, but have an average length of 14 kb. DoGs are retained in the nucleus and largely remain localized close to their site of transcription (Vilborg et al. 2015; Vilborg et al. 2017; Hennig et al. 2018).

Upon influenza and HSV-1 infection, the induction of readthrough transcripts is orchestrated in part by viral proteins NS1 and ICP27, respectively. These proteins interfere with transcription termination by interacting with host cleavage and polyadenylation factors (Nemeroff et al. 1998; Bauer et al. 2018; Wang et al. 2020). Accordingly, knockdown of the catalytic subunit of the cleavage and polyadenylation apparatus, CPSF73, leads to a partial induction of DoGs (Vilborg et al. 2015). However, the host-dependent mechanisms underlying the induction of DoGs upon environmental stress, such as hyperosmotic stress, remain unknown.

Readthrough transcripts have been previously characterized using nuclear RNA sequencing, 4sU-sequencing, mNET-seq and total RNA sequencing (Vilborg et al. 2015; Rutkowski et al. 2015; Hennig et al. 2018; Bauer et al. 2018; Grosso et al. 2015). While these techniques enable the genome-wide identification of DoG transcripts, most do not provide quantitative information regarding the nascent transcription profiles that coincide with DoG induction. Here we use Transient Transcriptome (TT) sequencing coupled with TimeLapse (TL) chemistry (Schwalb et al. 2016; Schofield et al. 2018) to characterize the transcriptional profiles that accompany DoG induction upon hyperosmotic stress. This technique enables the reliable detection of nascent RNAs by using short pulses of metabolic labeling coupled with transcript enrichment and nucleoside recoding chemistry.

The Integrator complex is important for the 3’ end processing of various noncoding RNAs, such as small nuclear RNAs (snRNAs), enhancer RNAs, viral microRNAs, lncRNAs and replication-dependent histone genes (Albrecht and Wagner. 2012; Lai et al. 2015; Xie et al. 2015; Nojima et al. 2018; Skaar et al. 2015). This complex also regulates promoter-proximal Pol II pausing at protein-coding genes (Stadelmayer et al. 2014, Gardini et al. 2014; Elrod et al. 2019). Integrator subunit 11 (Int11) is the catalytic subunit of the complex and forms a heterodimer with Int9 (Baillat et al. 2005; Wu et al 2017). These subunits are analogous to CPSF73 and CPSF100 (Dominski et al. 2005; Baillat and Wagner. 2015), respectively, which are essential for the 3’ processing of mRNAs (Mandel et al. 2006; Shi et al. 2009).

Here, we report that hyperosmotic stress causes widespread transcriptional repression, from which a minority of human genes escape. Yet, we observe the induction of DoGs from genes that experience different transcriptional responses to hyperosmotic stress. ChIP-seq experiments demonstrate that Pol II binding profiles correlate with the transcriptional output of genes after hyperosmotic stress. Furthermore, our proteomics results reveal changes in the Pol II interactome after stress. Specifically, we observe a decreased interaction between Pol II and subunits of the Integrator complex, while the interactions between Pol II and factors known to mediate the termination of protein-coding genes are unchanged. Correspondingly, siRNA knockdown of Int11 was sufficient to induce readthrough transcription at hundreds of genes. The set of genes that produce readthrough transcripts upon depletion of functional Int11 partially overlaps the collection of genes that give rise to DoGs after hyperosmotic stress. Together, our results provide mechanistic insight into the biogenesis of a recently characterized class of noncoding RNAs and reveal a novel role for the Integrator complex in transcriptional regulation.

## Results

### Hyperosmotic stress causes widespread transcriptional repression

We investigated the nascent transcriptional profiles accompanying DoG induction by performing TT-TL-seq (Schofield et al. 2018) of untreated HEK-293T cells and cells exposed to hyperosmotic stress (Figure 1A). Specifically, we treated cells with 80 mM KCl for a total time course of sixty minutes. For the last five minutes of KCl treatment, we added the nucleoside analog 4-thiouridine (s^4^U) in order to label RNAs that were being actively transcribed (Schwalb et al. 2016). After extracting RNA from HEK-293T cells, RNA from *Drosophila* S2 cells was added to each sample as a normalization control to ensure accurate differential expression analysis (Lovén et al. 2012; Chen et al. 2015). RNAs containing s^4^U were then biotinylated using MTS chemistry and enriched on streptavidin beads (Duffy et al. 2015). Finally, U-to-C mutations were induced using TimeLapse chemistry in order to reliably assess the nascent nature of the enriched RNAs (Schofield et al. 2018). TT-TL-seq experiments were performed using conditions that were previously found to induce DoGs (Vilborg et al. 2017). RT-qPCR results from HEK-293T cells exposed to hyperosmotic stress for different time points confirmed that DoGs were robustly induced in these cells after 1 hour of treatment (Figure S1A), consistent with previous results from NIH-3T3 cells (Vilborg et al. 2017).

**Figure 1:**
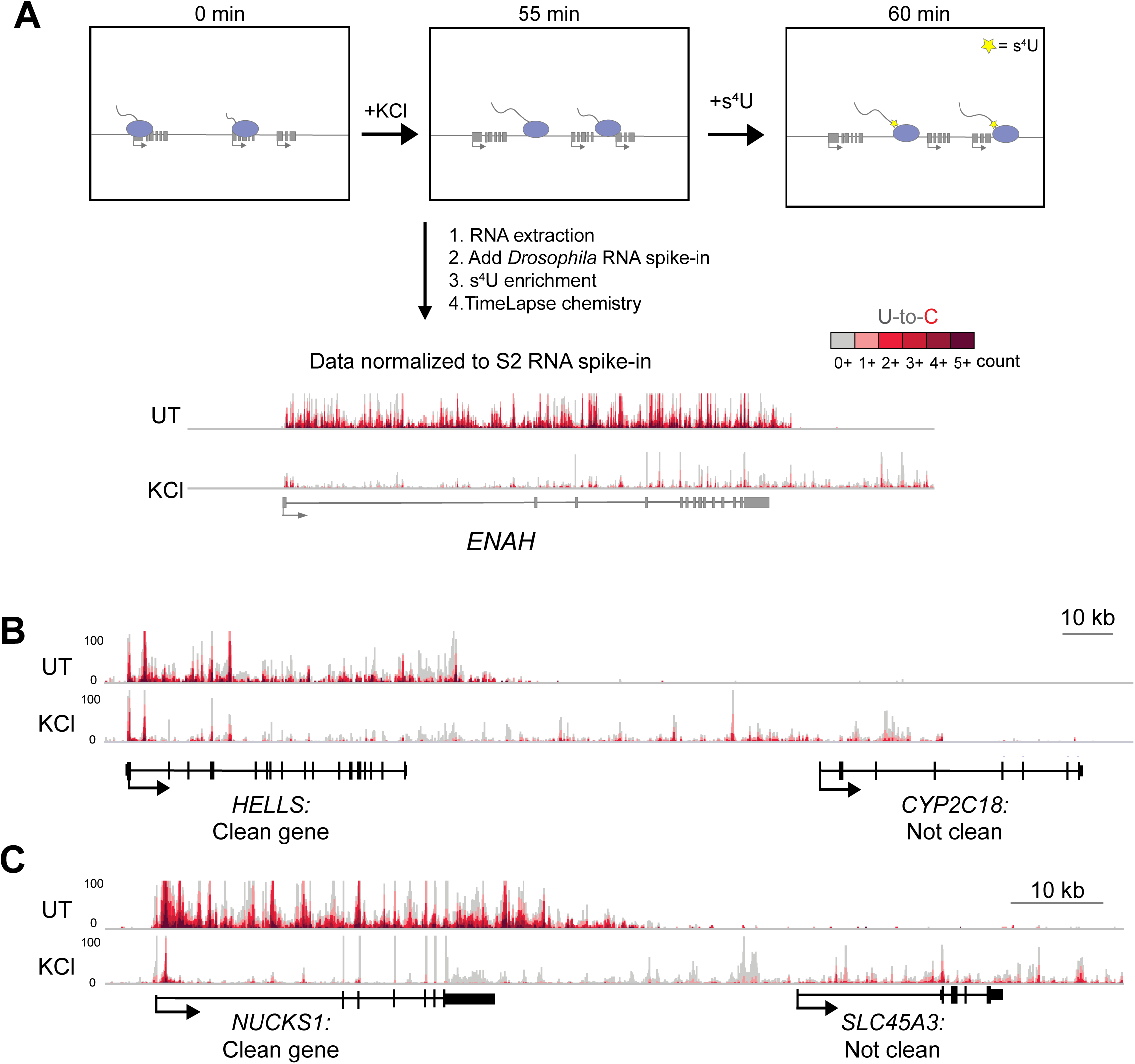
TT-TL-seq reveals transcriptional profiles that accompany DoG induction after hyperosmotic stress. A) Experimental setup for TT-TimeLapse sequencing (TT-TL-seq) experiments in HEK-293T cells. An arrow indicating directionality marks the beginning of each transcription unit. Exons are shown as rectangles and Pol II molecules are light purple ovals with attached nascent RNAs. Genome browser view of TT-TL-seq data for *ENAH* provide an example of results from untreated (UT) and KCl-treated (KCl) cells after normalization to the spike-in control. B) Browser image of TT-TL-seq data exemplifying a clean gene (*HELLS*) and a gene that does not meet the criteria for a clean gene (*CYP2C18*). The DoG produced from HELLS reads into CYP2C18 and makes it seem as if the latter is transcriptionally activated by hyperosmotic stress (log_2_ fold change = 5.39). C) The DoG produced from *NUCKS1* is assigned to *SLC45A3* because of extensive read-in transcription. Here, read-in transcription also complicates accurate differential expression analysis for *SLC45A3* (log_2_ fold change = 4.13).

It has been previously observed that read-in transcription of DoGs into neighboring genes often leads to the mis-characterization of overlapping transcripts as being activated by stress conditions (Rutkowski et al. 2015; Hennig et al. 2018; Cardiello et al. 2018, Roth et al. 2020; Figure 1B). Additionally, read-in transcription confounds the assignment of DoGs to the corresponding parent gene (Figure 1C). Our analyses suggest that around 55% of expressed genes experience read-in transcription after hyperosmotic stress (Table S1). Therefore, to ensure accurate differential expression analyses and DoG characterization, we generated a sub-list of genes, which we refer to as “clean genes” throughout the manuscript. The term “clean genes” describes genes that are expressed, do not overlap with DoGs from neighboring genes on either strand and have a higher expression within the gene body than the region 1kb upstream of the gene’s transcription start site (TSS) (Roth et al. 2020; Figure 1B & C). We identified 4584 clean genes in HEK-293T cells after hyperosmotic stress and analyzed their transcriptional regulation (Table S1).

Consistent with previous reports that analyzed steady-state RNAs in human cells (Amat et al. 2019), we were able to identify changes in transcriptional responses after hyperosmotic stress. Our results reveal predominantly decreases in nascent transcript levels after stress (Figure 2A). Specifically, we observe a 3-fold decrease in the number of normalized read counts corresponding to clean gene bodies after KCl treatment (Figure S1D). However, we find that a subset of clean genes is able to bypass this transcriptional repression (Figures 2B, C & S1E), including *GADD45B* (Figure 2D), which is known to be induced by hyperosmotic stress (Mak and Kültz 2004). More than 88% of clean genes were repressed after hyperosmotic stress, while only 3% of clean genes were upregulated (Figures 2C & S1F). To validate these observations, we extracted total RNA from untreated and KCl-treated cells and performed RT-qPCR using primers targeting intronic regions of several repressed genes. Results from HEK-293T cells and from SK-N-BE(2)C cells confirm the transcriptional decrease of three representative genes upon hyperosmotic stress (Figure S1B & C). RT-qPCR results from both cell lines also demonstrate that mature mRNA levels for the three genes remain unaffected, while levels of readthrough transcripts increase after hyperosmotic stress. Together our results reveal that hyperosmotic stress alters the nascent transcription profiles of cells by repressing thousands of genes and activating a small subset of genes.

**Figure 2:**
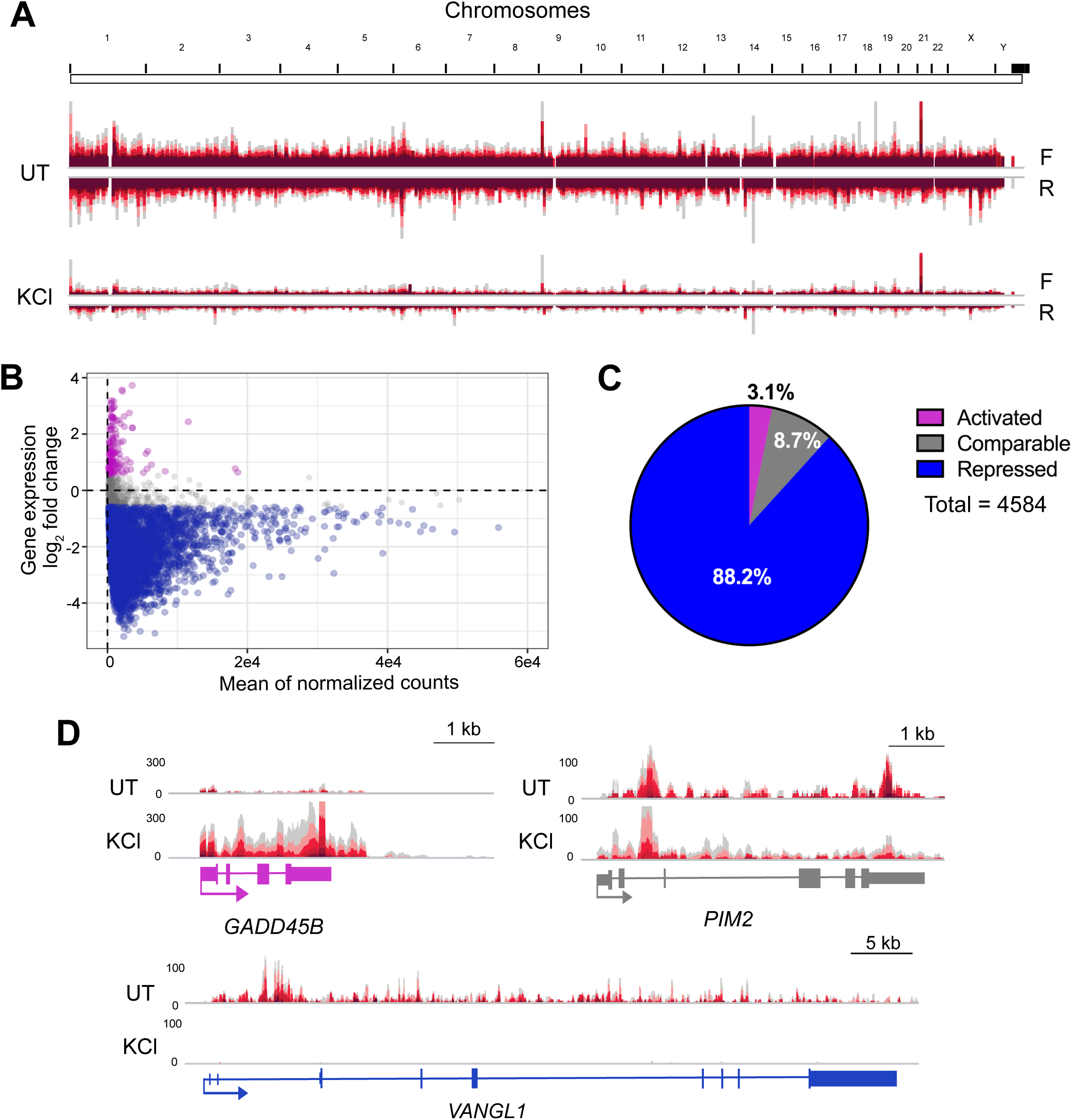
Hyperosmotic stress leads to widespread transcriptional repression. A) Whole-genome view of TT-TL-seq normalized reads for forward (F) and reverse (R) strands in UT and KCl samples. B) Minus average plot showing the log_2_ fold change for clean genes on the y-axis and the mean of normalized counts on the x-axis. Activated genes are shown in purple, genes retaining comparable expression levels are shown in gray and repressed genes are shown in blue (n=4584). C) Pie-chart illustrating the percentage of clean genes within each of the 3 categories of transcriptional regulation (activated gene log_2_ FC > 0.58; comparable gene log_2_ FC < 0.58 but > −0.58; repressed gene log_2_ FC < −0.58). D) Brower shots of TT-TL-seq tracks from HEK-293T cells for *VANGL1*, which is transcriptionally repressed by hyperosmotic stress (log_2_ FC = −4.97), *PIM2*, which retains comparable expression levels after KCl treatment (log_2_ FC = 0.28), and *GADD45B*, which is activated by hyperosmotic stress (log_2_ FC = 3.72).

### Stress-induced readthrough transcripts arise independent of gene transcription levels

Consistent with the observed widespread transcriptional repression, normalized read counts for nascent RNAs (TT-TL-seq) within the bodies of DoG-producing clean genes decreased after hyperosmotic stress, while read counts corresponding to DoG regions increased (Figure 3A). However, log_2_ fold changes in nascent RNAs of DoG-producing clean genes show that DoGs are produced from genes that experience all three types of transcriptional responses (Figure 3B; Table S1). These results demonstrate that 2.9% of DoGs arise from activated clean genes, 87.8% of DoGs arise from repressed clean genes and 9.3% of DoGs arise from clean genes that retain comparable levels of expression in stressed and unstressed HEK-293T cells (Figure 3B). We then asked whether DoGs preferentially arise from genes that are transcriptionally repressed upon hyperosmotic stress. Interestingly, the percentage of DoG-producing genes within each class of transcriptional regulation is consistent, comprising 12-14% (Figure 3C). We conclude that DoGs are produced regardless of the transcriptional level of their upstream genes (Figure 3D).

**Figure 3:**
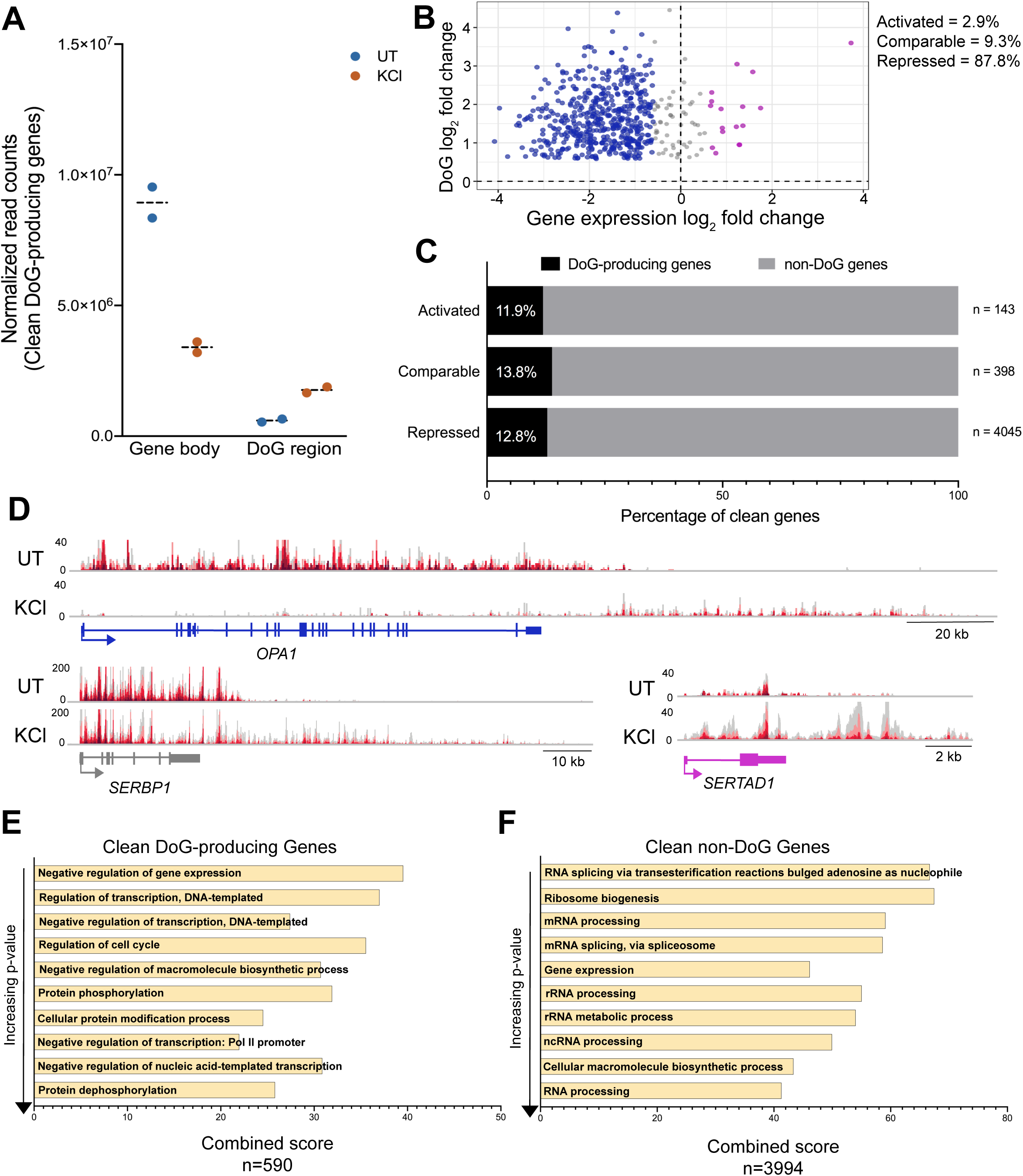
DoGs arise regardless of the transcriptional levels of their upstream genes upon hyperosmotic stress. A) Interleaved scatter plot showing the sum of normalized TT-TL-seq read counts of DoG-producing clean genes and corresponding DoG regions (n=590) in untreated and KCl-treated HEK-293T cells for two biological replicates. B) Scatter plot showing clean gene log_2_ fold change (FC) for the gene body on the x-axis and the log_2_ FC for the DoG region on the y-axis. DoG-producing genes that are transcriptionally activated upon stress are represented in purple, genes retaining comparable levels of expression in gray and genes that are repressed in blue. C) Bar graph showing the percentage of DoG-producing clean genes (black) within each category of transcriptional regulation. D) Browser image showing UT and KCl TT-TL-seq reads for *OPA1*, a transcriptionally repressed DoG-producing clean gene (gene log_2_ FC = −3.22), for *SERBP1*, which retains comparable expression levels after stress (gene log_2_ FC = −0.45) and for a transcriptionally activated DoG-producing clean gene, *SERTAD1* (gene log_2_ FC = 1.23). E, F) Bar graphs show gene ontology combined scores for the ten most significantly enriched biological processes in order of increasing p-value for E) DoG-producing clean genes and for F) non-DoG clean genes. Combined scores are the product of the p-value and the z-score as calculated by Enrichr (Chen et al. 2013).

### Clean DoG-producing genes are functionally enriched for transcriptional repression

Previous analyses of DoG-producing genes did not reveal any functional enrichment (Vilborg et al. 2017). We suspected that the challenge of assigning DoGs to the correct gene of origin because of their extension into neighboring genes may have complicated previous analyses (Figure 1C). Therefore, we revisited the question of whether DoG-producing genes are enriched for certain biological processes using only clean genes. We performed gene ontology analysis of DoG-producing clean genes and clean genes that fail to generate DoGs (non-DoG genes) using Enrichr (Chen et al. 2013; Kuleshov et al. 2016; Table S2). Interestingly, five out of the ten enriched terms with the most significant p-values for DoG-producing genes are related to transcriptional repression (Figure 3E). The remaining five terms are related to transcriptional regulation and protein modifications. Non-DoG genes do not show such a striking enrichment for terms related to transcriptional repression compared to other terms (Figure 3F). Instead, these genes are strongly enriched for general processes related to RNA processing.

### Pol II is redistributed along the genome after hyperosmotic stress

We used anti-Pol II ChIP-seq to investigate how Pol II binding correlates with the transcriptional profiles observed through TT-TL-seq in untreated HEK-293T cells and in cells exposed to hyperosmotic stress. Meta-analyses of DoG-producing genes confirmed a redistribution of Pol II molecules to the downstream regions in stressed cells (Figures 4A & S4A), consistent with DoG transcription and with previous observations (Cardiello et al. 2018; Heinz et al. 2018). As expected, this redistribution of Pol II molecules was not observed downstream of non-DoG genes (Figures 4A & S4A). We asked to what extent the redistribution of Pol II molecules into the regions downstream of genes was responsible for the widespread transcriptional repression observed by TT-TL-seq. Alternatively, the decrease in transcription might be caused by a decrease in Pol II binding at repressed genes. It has been previously reported that hyperosmotic stress induced by NaCl decreases Pol II binding to DNA (Amat et al. 2019). Our results show a similar number of Pol II binding peaks after KCl treatment (Figure S4B). We observe that a subset of overlapping peaks show increased Pol II binding after stress, while the majority of the identified peaks either retain comparable levels of Pol II binding or exhibit reduced Pol II binding (Figure S4C & D). Consistently, we found that Pol II binding decreases over most clean gene bodies after KCl treatment (Figure 4B).

**Figure 4:**
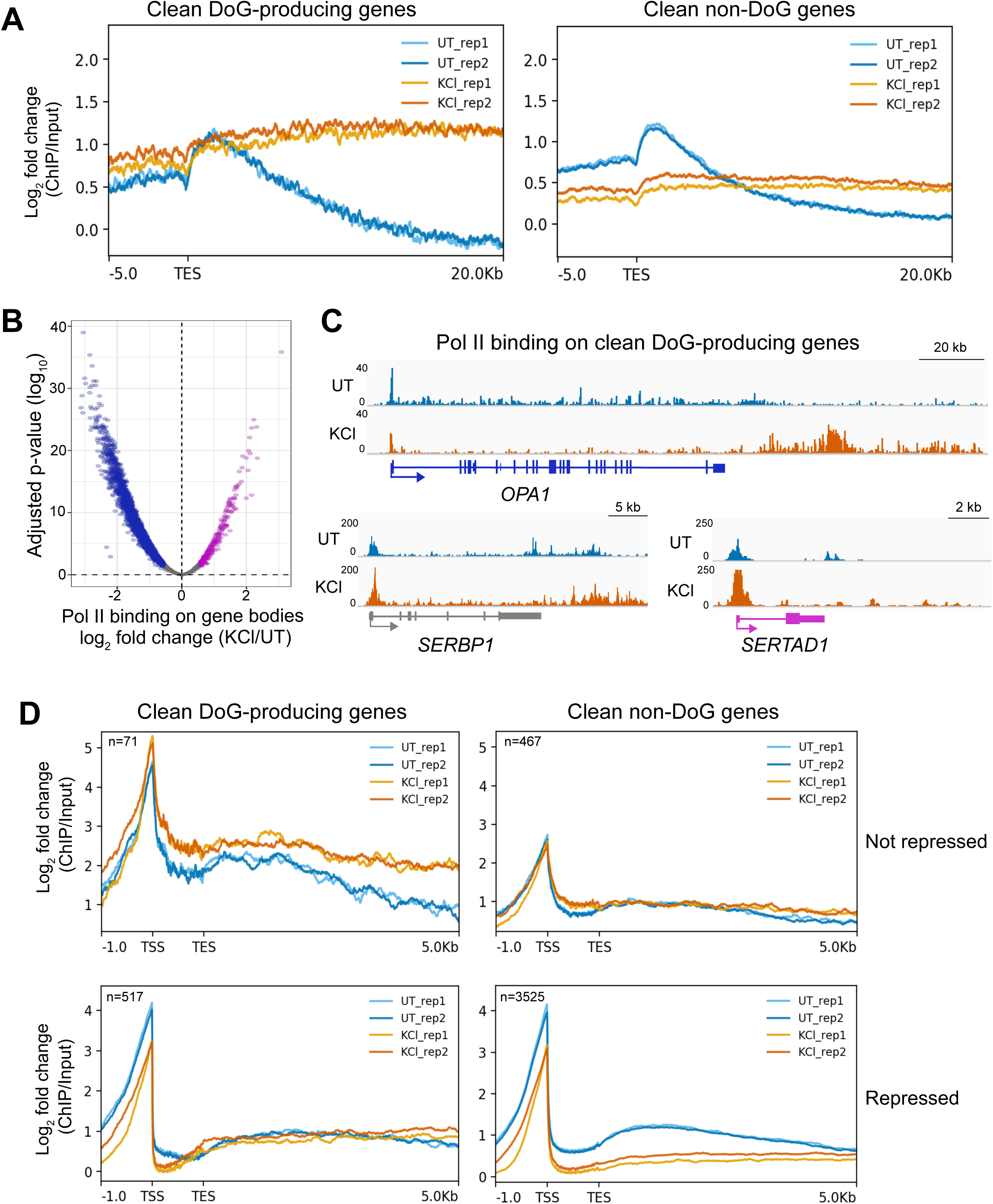
Hyperosmotic stress causes a redistribution of Pol II molecules across the genome. A) Meta plots showing the log_2_ fold change of anti-Pol II ChIP-seq data normalized to input across the annotated transcription end sites (TES) of DoG-producing clean genes (n=590) and of clean non-DoG genes (n=3994) from UT and KCl-treated HEK-293T cells. B) Volcano plot showing the log_2_ fold change of read counts from anti-Pol II ChIP-seq data normalized to input for clean genes on the x-axis and the corresponding log_10_ adjusted p-values on the y-axis. Genes with increased Pol II binding across the gene body are shown in purple, genes retaining comparable levels of Pol II binding are shown in gray and those with decreased Pol II binding across the gene body are shown in blue. C) Browser images of DoG-producing clean genes showing anti-Pol II ChIP-seq tracks normalized to input: *OPA1* is a repressed gene, *SERBP1* retains comparable expression levels and *SERTAD1* is activated by hyperosmotic stress according to TT-TL-seq data. D) Meta plots showing Pol II binding in DoG-producing clean genes and clean non-DoG genes that are not repressed (top) and genes that are repressed (bottom). Two biological replicates are shown for each meta plot for UT and KCl samples.

We verified Pol II occupancy patterns for representative genes within each category of transcriptional regulation. Pol II binding profiles (Figure 4C) of *OPA1*, a DoG-producing gene that is transcriptionally repressed, show decreased binding close to the transcription start site and a shift of Pol II peaks into the downstream region in stressed cells. On the other hand, for *SERTAD1*, a DoG-producing gene that is activated by hyperosmotic stress, and for *SERBP1*, which retains comparable levels of expression according to our TT-TL-seq data, we observe an increase in Pol II binding close to each TSS that extends downstream of these genes. Finally, we assessed the distribution of Pol II molecules across repressed clean genes and across clean genes that are not repressed, i.e. genes that are either activated or retain comparable levels of expression after hyperosmotic stress. Consistent with the individual gene examples discussed above, clean DoG-producing genes that are not repressed by hyperosmotic stress show increased Pol II binding close to their TSSs and across the gene body, while genes that are transcriptionally repressed by hyperosmotic stress show decreased levels of Pol II binding across the gene body. Similar effects were also observed for clean non-DoG genes (Figure 4D). Our results are, therefore, in agreement with the nascent transcription profiles derived from our TT-TL-seq data (Figures 2 & 3) and support a model where hyperosmotic stress induces both a redistribution of Pol II molecules along the genome and a decrease in Pol II binding at repressed genes.

### The Pol II interactome is altered by hyperosmotic stress

To gain insight into the mechanisms that promote DoG formation upon hyperosmotic stress, we performed immunoprecipitation of chromatin-bound Pol II coupled with mass spectrometry analysis (Harlen et al. 2016) in SK-N-BE(2)C cells (Figure 5A). This human neuroblastoma cell line, where KCl-induced DoGs were originally described (Vilborg et al. 2015), was used because we observed more robust DoG induction in these cells than in HEK-293T cells (Figure S1B & C). We confirmed efficient isolation of chromatin fractions through western blots (Figure S5A). Overall, our results reveal distinct changes in the Pol II interactome (Figure 5B). Peptides corresponding to several subunits of the cleavage and polyadenylation apparatus, as well as to Xrn2 were detected in both untreated and KCl-treated samples (Figure 5C) at least once across seven sets of experiments. The presence of Xrn2 and CPSF1 in the anti-Pol II immunoprecipitates of untreated and KCl-treated cells was validated through western blots (Figure S5B-D). In contrast, 11 of the 14 subunits of the Integrator complex were not detected among the Pol II interactors in the KCl-treated samples (Table S3). To validate this observation, we performed western blots to detect Pol II in chromatin-bound anti-Integrator subunit 3 (Int3) immunoprecipitates from SK-N-BE(2)C and HEK-293T cells. These results confirmed decreased binding between Pol II and Int3 in KCl-treated samples compared to untreated samples from both cell lines (Figure 5D & E). To ensure that the decreased detection of Integrator subunits was not due to protein degradation, we also assessed the levels of Int3 and Int11 in whole cell lysates from SK-N-BE(2)C cells. Our western blots confirmed that the levels of these Integrator subunits do not decrease in cells after hyperosmotic stress (Figure S5E).

**Figure 5:**
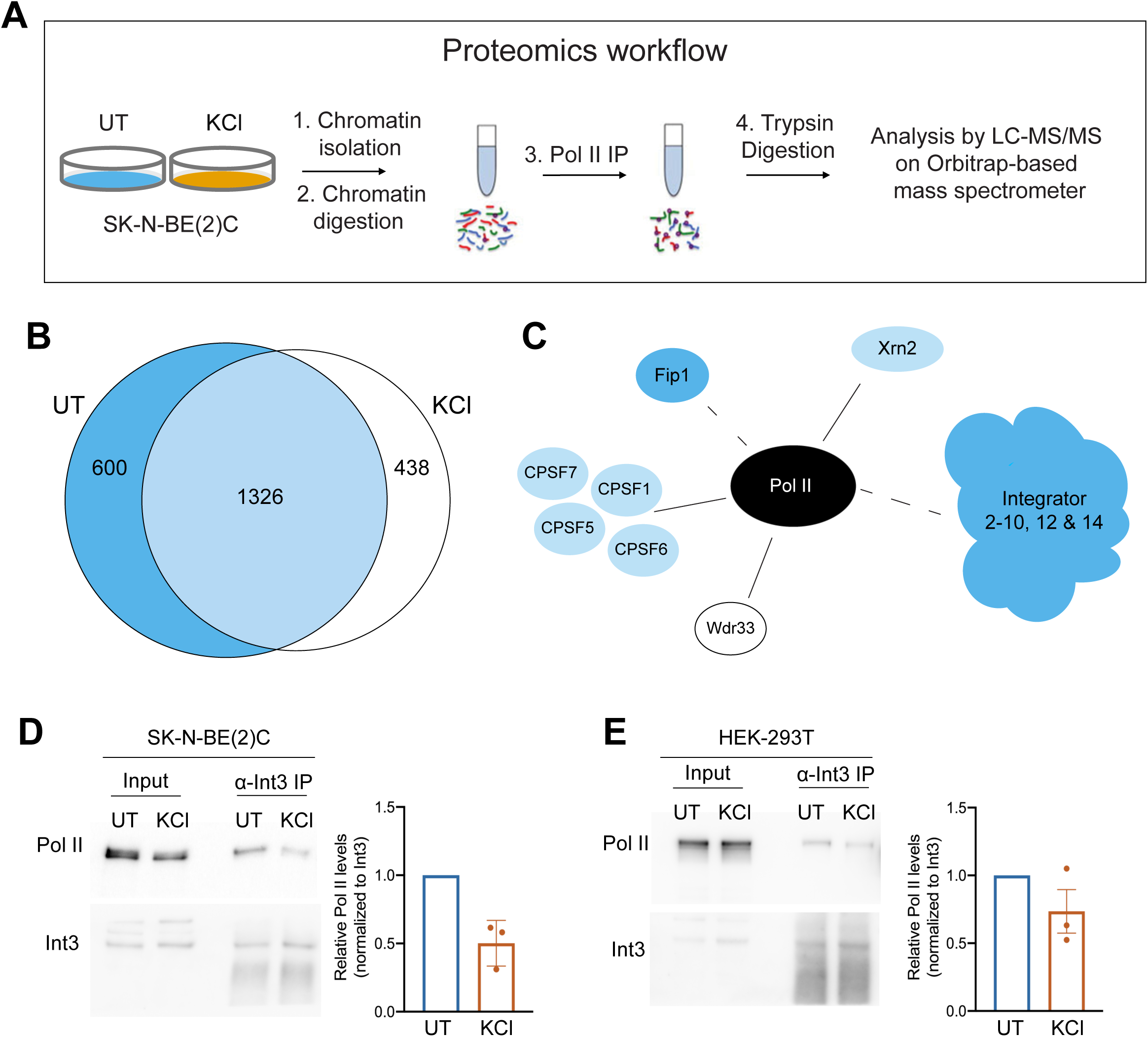
The Pol II interactome changes upon hyperosmotic stress. A) Experimental setup for mass spectrometry of chromatin-bound anti-Pol II immunoprecipitates from SK-N-BE(2)C cells. B) Overlap between proteins found in UT and KCl-treated samples (1326). 600 proteins were found only in the UT samples (blue) and 438 proteins were found only in the KCl-treated samples at least once across seven replicates (white). C) Cartoon depiction of changes in Pol II interactions with termination factors. Solid lines represent interactions that remain after hyperosmotic stress, while dashed lines represent interactions found only in the untreated samples. Proteins detected only in the untreated sample are shown in blue, proteins present only in the KCl-treated sample are shown in white and proteins present in both samples are shown in light blue, as in the Venn diagram shown in 5B. D, E) Western blots of chromatin-bound anti-Int3 immunoprecipitates reveal decreased levels of associated Pol II in KCl-treated samples compared to untreated samples from D) SK-N-BE(2)C and from E) HEK-293T cells. Chromatin fractions from these cell lines were used as input.

Together, our proteomic results show that hyperosmotic stress induces changes to the Pol II interactome that could contribute to the transcriptional landscape revealed by high throughput sequencing. Specifically, we observed that important termination factors interact with RNA Pol II despite cellular stress, while the interaction between Pol II and the Integrator complex decreases after stress.

### Depletion of Integrator endonuclease subunit leads to DoG production

The Integrator complex regulates transcription termination at many noncoding RNA loci and has been shown to bind the 3’ end of certain protein-coding genes (reviewed in Baillat and Wagner 2015; Gardini et al. 2014). Since the interaction between Pol II and Integrator decreases after hyperosmotic stress (Figure 5), we investigated whether knocking down the catalytic subunit of the complex, Int11, using siRNAs was sufficient to induce DoGs. We transfected HEK-293T cells with an siRNA against Int11 (siInt11) or with a non-targeting siRNA control (siC). HEK-293T cells stably expressing FLAG-tagged, siRNA-resistant wild-type (WT) or catalytically inactive Int11 (E203Q) were also transfected with siInt11 (Baillat et al. 2005; Xie et al. 2015) for 72 hours. DoG levels were then measured through RT-qPCR. Results obtained using DoG-specific primers demonstrate that knockdown of endogenous Int11 is sufficient to induce DoGs. Moreover, expression of exogenous WT Int11 reduces DoG induction compared to samples expressing the E203Q mutant or to samples expressing no rescue (Figure 6A).

**Figure 6:**
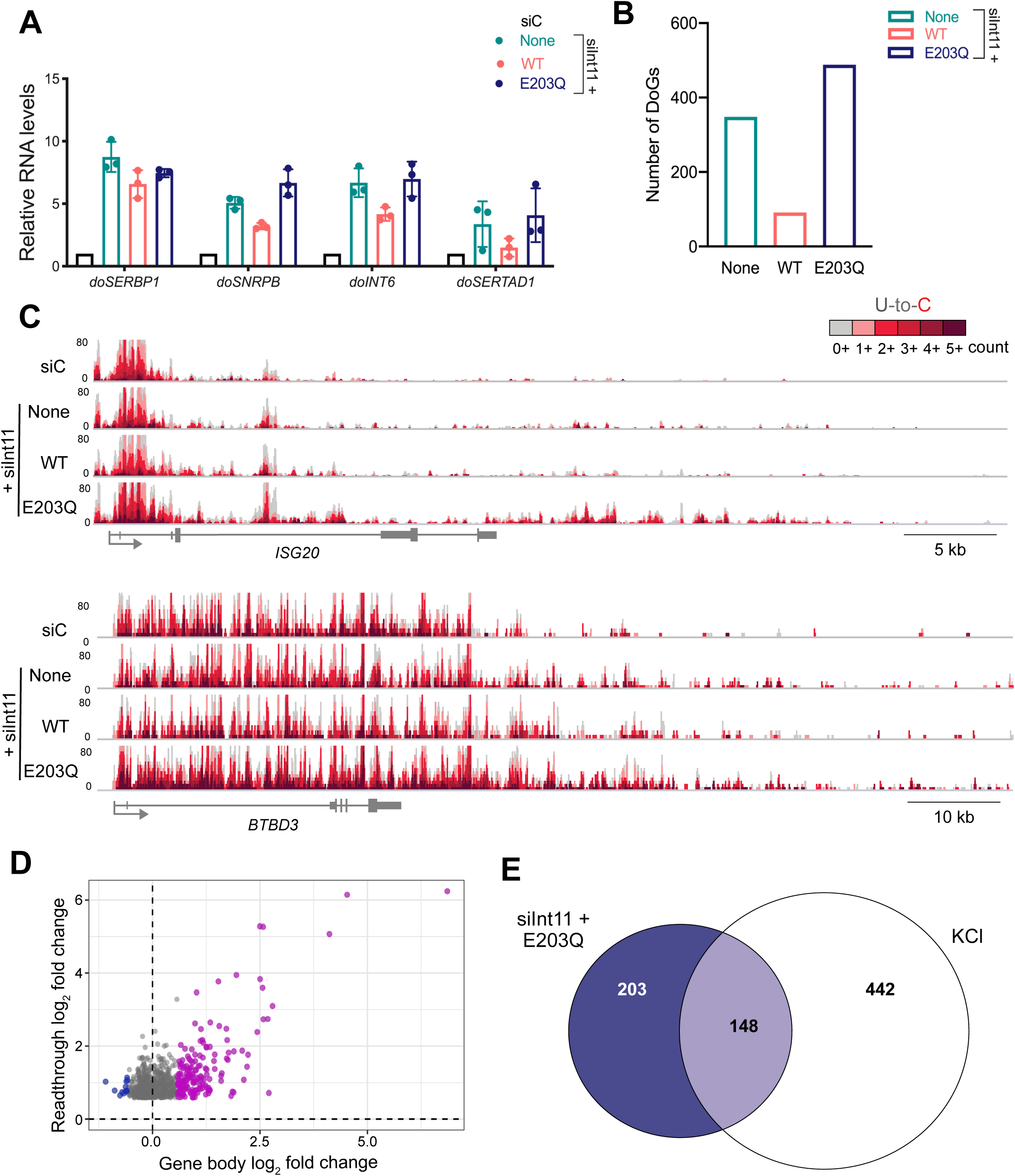
Depletion of Integrator nuclease subunit leads to readthrough transcription. A) RT-qPCR analysis of DoG levels in HEK-293T cells transfected with a scrambled siRNA control (siC) or an siRNA targeting endogenous Int11 (siInt11). The siInt11-transfected cells expressed either exogenous wild-type Int11 (WT), catalytically inactive Int11 (E203Q) or no exogenous Int11 (None). B) Bar graph showing the number of DoGs induced after siRNA knockdown of endogenous Int11. C) Browser image of TT-TL-seq data for two genes that produce DoGs upon depletion of functional Int11. D) Scatter plot showing gene expression log_2_ fold change (FC) for siInt11 + E203Q HEK-293T cells on the x-axis and the log_2_ FC of the corresponding readthrough transcripts on the y-axis. Genes that are activated in siInt11 + E203Q cells compared to siC-transfected cells are shown in purple, genes that are unaffected are gray and genes that are repressed are blue. All readthrough sites identified in the siInt11 + E203Q sample are represented in this plot (n=840). E) Venn diagram displaying overlap between the identities of clean genes that produce readthrough transcripts in siInt11 + E203Q cells (dark blue) and those of DoG-producing clean genes in KCl-treated samples (white).

Endogenous Int11 protein levels were reduced by ∼90% in siInt11-transfected samples (Figure S6A). The efficiencies of exogenous WT and E203Q mutant Int11 expression were verified by measuring the levels of unprocessed U2 through RT-qPCR (Skaar et al. 2015; Figure S6B): unprocessed U2 accumulates in cells without rescue (None) or expressing the E203Q mutant Int11, but not in cells transfected with siC or cells expressing exogenous WT Int11. Furthermore, northern blots using probes against snRNA U2 showed that levels of mature U2 remained unaffected in our knockdown samples (Figure S6C), consistent with previous reports (Tatomer et al. 2019). This argues against the possibility that the effects of Int11 knockdown on DoG production are because of the role of Integrator in snRNA processing.

We assessed the extent to which knockdown of endogenous Int11 induces DoGs genome-wide. To increase cell viability, we performed TT-TL-seq on HEK-293T cells transfected with an siRNA against Int11 for 48 hours (Figure S6D). Results obtained from cells lacking functional Int11 reveal hundreds of readthrough sites across the genome that are induced by more than 1.5-fold compared to the siC-transfected sample (Figure 6B; Table S4). According to TT-TL-seq data, induction of readthrough transcription after depletion of endogenous Int11 was most evident upon expression of the E203Q mutant (Figure 6B & C). We found that readthrough transcripts observed in siInt11-transfected cells expressing no rescue and in cells expressing the E203Q mutant Int11 are highly correlated (Figure S6E). As expected, the most highly induced sites of readthrough transcription (Table S4) corresponded to snRNA genes (Baillat et al. 2005; Albrecht and Wagner. 2012). We were also able to detect readthrough downstream of lncRNA and histone genes as previously described (Nojima et al. 2018; Skaar et al. 2015). However, most of the identified readthrough sites were downstream of protein-coding genes (Table S4). Out of the 840 readthrough transcripts induced by knockdown of functional Int11 (E203Q sample), 489 had more than 80% read coverage in the region 5 kb downstream of the annotated termination site of the gene of origin and, therefore, met all criteria to be classified as DoG RNAs (Figure 6B & C; Table S4).

Depletion of Integrator subunits has been shown to alter the transcriptional levels of certain genes (Gardini et al. 2014; Elrod et al. 2019; Tatomer et al. 2019). Consistently, we found that depletion of functional Int11 in HEK-293T cells differentially affects more than a thousand genes (Figure S6F). The expression levels of the parent genes upstream of readthrough transcripts induced upon depletion of functional Int11 reveals that readthrough transcripts predominantly arise from upregulated genes or from genes that retain comparable levels of expression after knockdown, while very few arise from genes that are transcriptionally repressed (Figure 6D).

Given our observation that the interaction between subunits of the Integrator complex and Pol II are disrupted by hyperosmotic stress, we asked how the identities of genes producing readthrough transcripts upon depletion of functional Int11 compare to genes that produce DoGs after hyperosmotic stress. We identified 232 clean genes that produce readthrough transcripts in siInt11-transfected cells and 351 clean genes that produce readthrough transcripts in the siInt11 + E203Q mutant sample. Comparison with the 590 DoG-producing clean genes identified in cells treated with hyperosmotic stress showed that up to 25% of KCl-induced DoGs are detected at loci that also produce readthrough transcripts after depletion of functional Int11 (Figure 6E & S6G). These readthrough transcripts are generally more robustly induced after KCl treatment than upon depletion of functional Int11 (Figure S6H), suggesting that, although Int11 knockdown is sufficient to produce readthrough transcription, decreased interactions between Integrator and Pol II are not solely responsible for DoG induction upon hyperosmotic stress.

## Discussion

In this study we have used TT-TimeLapse sequencing to evaluate nascent transcription profiles that accompany DoG induction under conditions of hyperosmotic stress (Figure 1). We observe that hyperosmotic stress triggers transcriptional repression that is widespread and of greater magnitude than anticipated (Robbins et al. 1970; Amat et al. 2019). Our results reveal that after an hour-long exposure to mild conditions of hyperosmotic stress, only ∼12% of clean genes escape this transcriptional repression (Figure 2). Interestingly, we find that DoG RNAs are produced independent of the transcriptional levels of their upstream genes and that DoG-producing genes are functionally enriched for transcriptional repression according to gene ontology analyses (Figure 3). Pol II binding profiles demonstrate a correlation with nascent transcription profiles across DoG-producing clean genes and non-DoG genes (Figure 4). Additionally, proteomic analysis of the Pol II interactome revealed decreased binding between the Integrator complex and Pol II upon exposure to hyperosmotic stress (Figure 5). Depletion of the catalytic subunit of the Integrator complex, Int11, was sufficient to induce readthrough transcription downstream of hundreds of loci. Genes that produce readthrough transcripts upon depletion of functional Int11 and genes that produce DoGs upon hyperosmotic stress partially overlap (Figure 6).

Our analysis of clean genes has enabled us to accurately characterize the relationship between DoGs and their upstream transcripts (Figures 1 & 3). Consistent with previous observations in heat shock-treated cells (Cardiello et al. 2018), we find that DoGs can be produced from genes that experience different transcriptional responses to hyperosmotic stress. Importantly, a similar percentage of clean genes that are activated, repressed or unchanged are able to produce DoGs after hyperosmotic stress, revealing that transcriptional levels and DoG production are not mechanistically linked. Furthermore, our analyses suggested that around 55% of genes that are expressed in HEK-293T cells might be affected by read-in transcription upon exposure to hyperosmotic stress (Table S1). Intriguingly, Rutkowski and colleagues have previously reported that read-in transcription from upstream genes negatively affects the splicing of downstream genes upon HSV-1 infection (Rutkowski et al. 2015). Similarly, Shalgi and colleagues reported widespread disruptions in co-transcriptional splicing upon heat shock (Shalgi et al. 2014). Reports from Hennig and colleagues confirm aberrant splicing events upon heat shock and hyperosmotic stress in human foreskin fibroblast cells (Hennig et al. 2018). All these findings suggest that altered splicing patterns may be a consequence of DoG production and highlight the extensive impact of DoG production on the transcriptome of stressed cells. Investigating the diversity of unprocessed transcripts that accumulate upon stress will provide a better understanding of how transcriptional and processing dynamics are impacted during the cellular stress response.

Upon revisiting the question of whether hyperosmotic stress-induced DoG-producing genes show any functional enrichment, we found that many clean genes encode transcriptional repressors (Figure 3; Table S2). Transcriptional repression has also been reported under other DoG-inducing conditions, including heat shock and viral infections (Mahat et al. 2016; Rutkowski et al. 2015; Bauer et al. 2018). In light of this finding, it is tempting to speculate that DoGs could serve as a store of unprocessed transcripts that facilitate the cell’s survival upon prolonged stress by providing a source of important mRNAs that would require processing, but not active transcription, thereby reducing the cell’s energetic costs. DoG induction might further promote transcriptional repression during hyperosmotic stress by causing transcriptional interference at neighboring genes (Mazo et al. 2007; Muniz et al. 2017). Other terms of biological enrichment for DoG-producing genes were related to protein modifications, which are crucial for the rapid regulation of cellular processes. A repository of transcripts that encode these proteins might facilitate recovery from stress. Alternatively, given the transcriptional changes observed upon hyperosmotic stress, it is possible that DoGs serve to alter the chromatin landscape surrounding their parent and neighboring genes in order to facilitate their prolonged regulation throughout the stress response and recovery from stress.

In agreement with previous observations in heat shock (Cardiello et al. 2018), anti-Pol II ChIP-seq data from cells exposed to hyperosmotic stress demonstrate a decrease in Pol II binding across the bodies of repressed clean genes accompanied by a shift of elongating Pol II molecules into downstream regions (Figure 4). At loci that are not repressed by stress, we observe an increase in Pol II binding along the gene bodies that extends into their downstream regions. These observations are consistent with our TT-TL-seq results (Figures 2 & 3). The differing levels of Pol II occupancy observed across parent genes suggest an elegant and tightly regulated mechanism of transcriptional termination at DoG-producing genes.

Surprisingly, rather than observing decreased interactions between Pol II and components of the cleavage and polyadenylation apparatus, our proteomic analyses revealed that the binding between Integrator subunits and Pol II decreases after stress (Figure 5). We then asked whether depleting Int11, the catalytic subunit of the complex, using siRNAs is sufficient to induce DoGs. TT-TL-seq of siRNA-treated samples revealed that hundreds of DoGs are induced after functional Int11 is depleted from cells (Figure 6). Moreover, our results reveal that expression of wild-type Int11 reduces the number of DoGs induced in cells where endogenous Int11 has been knocked down compared to cells expressing no rescue. Additionally, we observe an increase in the number of DoGs induced upon expression of the E203Q mutant Int11. This argues that the nuclease activity of Int11 is important for repressing DoG production under homeostasis. These results highlight the importance of the Integrator complex in the regulation of mRNA production beyond the promoter proximal pause site (Elrod et al. 2019). It is possible that cells rely on the Integrator complex to mediate a mechanism of transcription termination at a subset of protein-coding genes that is an alternative to the canonical mechanism orchestrated by the cleavage and polyadenylation apparatus. This might explain why readthrough transcription is not observed downstream of all protein-coding genes. More work dissecting the role of the Integrator complex in mRNA 3’ end formation and transcription termination of protein-coding genes is imperative.

Our results reveal that as many as 25% of genes that produce DoGs upon KCl-treatment also produce readthrough transcripts upon depletion of functional Int11 (Figure 6). This partial overlap between Int11-depletion-dependent readthrough transcripts and KCl-induced DoGs suggests that the decreased binding of Integrator subunits to Pol II throughout the stress response partially contribute to DoG biogenesis in this context. Nonetheless, this comparison is limited by the temporal differences inherent to the experimental protocol. However, because of the changes imposed on the transcriptional landscape upon hyperosmotic stress (Figure 2), it is unlikely that the depletion of a single protein is enough to recapitulate the complexity of this response. Transcriptional regulation varies across these two conditions. Hyperosmotic stress induces widespread transcriptional repression to an extent that is not recapitulated upon depletion of functional Int11. Additionally, our results reveal that there are changes in Pol II binding patterns to DNA (Figure 4) and to other proteins (Figure 5; Table S3) after hyperosmotic stress. The full extent of DoG-induction upon cellular stress might also be supported by factors that have not yet been characterized. These could include a stress-induced increase in the processivity of elongating Pol II molecules, which would also result in readthrough transcription (Fong et al. 2015).

In agreement with the important roles of Xrn2 and CPSF73 in mediating the termination of protein-coding gene transcripts (Proudfoot. 2016), readthrough transcription is observed upon depletion of these proteins (Fong et al. 2015; Eaton et al. 2018; Eaton et al. 2020). However, unlike stress-induced readthrough transcription, the effect of depleting these proteins was not limited to a subset of genes. Using anti-Pol II immunoprecipitation of chromatin fragments coupled with mass spectrometry and western blots, we demonstrate that the overall binding of Pol II and important termination factors, including Xrn2 and subunits of the cleavage and polyadenylation apparatus, is not affected by hyperosmotic stress. Our data, however, do not exclude the possibility that important termination factors are redistributed along the genome after hyperosmotic stress. Anti-CPSF6 ChIP-seq from cells treated with NaCl to induce hyperosmotic stress revealed decreased binding of this subunit at a set of genes (Jalihal et al. 2019), yet it is unclear whether this was accompanied by a decrease in Pol II occupancy across the bodies of these genes. Moreover, anti-CPSF73 ChIP-qPCR data from cells exposed to heat shock suggest that recruitment of this factor decreases at activated DoG-producing genes (Cardiello et al. 2018). It is possible that at genes that are activated by stress the stoichiometry between Pol II and important termination factors is disrupted, which would contribute to DoG production. However, this model would not explain readthrough transcription at genes that retain comparable expression levels or genes that are transcriptionally repressed after stress, which comprise the majority of DoG-producing genes upon hyperosmotic stress (Figure 3B). These observations emphasize the need to further explore characteristic features of DoG-producing genes that might make them susceptible to failed termination despite the presence of important termination factors on chromatin.

## Acknowledgments

We thank Annsea Park, Kazimierz Tycowski, Salehe Ghasempur and other members of the Steitz lab for critical discussion of the manuscript. We are grateful to Samuel Roth for his assistance in troubleshooting our read-in analysis using ARTDeco. We thank Angela Miccinello for editorial assistance and Mingyi Xie and Mei-Di Shu for cells stably expressing exogenous Int11. This work was supported by a grant from the National Institutes of Health Common Fund 4D Nucleome Program (CA200147). N.A.R.M. is a Ford Foundation pre-doctoral fellow and J.A.S. is an investigator of the Howard Hughes Medical Institute.

## Author contributions

N.A.R.M. and J.A.S. conceived the project. All authors contributed to the experimental design. N.A.R.M. performed cellular experiments and ChIP-seq experiments. J.T.Z. performed enrichments, chemistry and library preparations for TT-TL-seq. N.A.R.M. and J.T.Z. did bioinformatic analyses of TT-TL-seq data. M.A. performed mass spectrometry and wrote the corresponding methods section. N.A.R.M. and J.A.S. wrote the rest of manuscript and prepared the figures with contributions from M.D.S., J.R., J.T.Z. and M.A.

---

**Figure S1:**
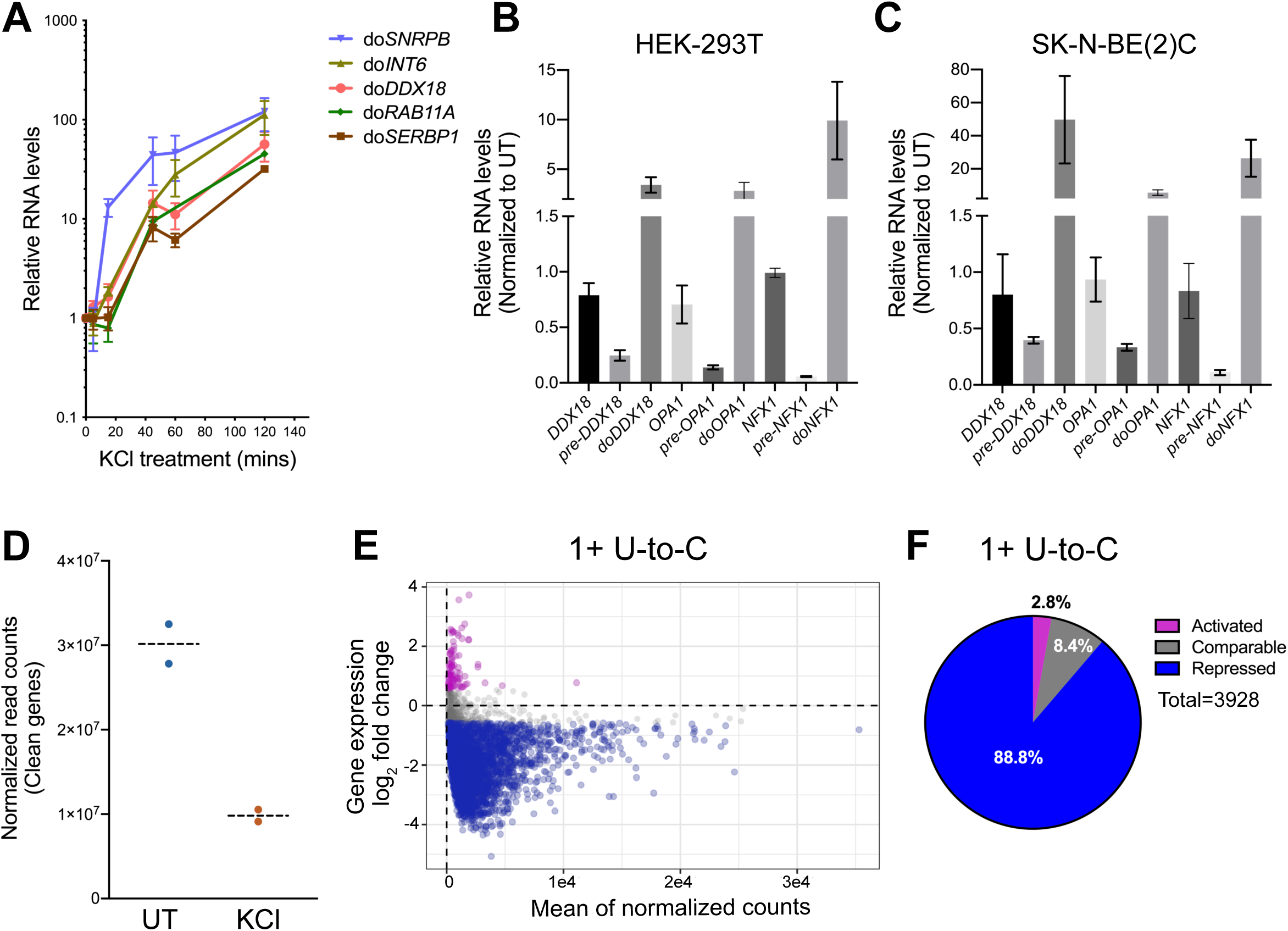
Hyperosmotic stress induces transcriptional repression. A) Time course of DoG induction from 5 DoG-producing genes in HEK-293T cells measured by RT-qPCR (n=3 for 5, 15, 45 and 60 minutes; n=2 for 120 minutes). B, C) Levels of mRNA, pre-mRNA and DoG RNA for *DDX18, OPA1* and *NFX1*, as determined by RT-qPCR in B) HEK-293T cells (n=3) compared to C) SK-N-BE(2)C cells (n=3). D) Interleaved scatter plot showing the sum of all normalized counts for clean genes in UT and KCl samples. E) Minus average plot showing reads with one or more U-to-C mutation induced by TimeLapse chemistry in untreated and KCl-treated samples. F) Pie chart illustrating the percentage of clean genes within each of the 3 categories of transcriptional regulation based on reads containing one or more U-to-C mutation induced by TimeLapse chemistry.

**Figure S4:**
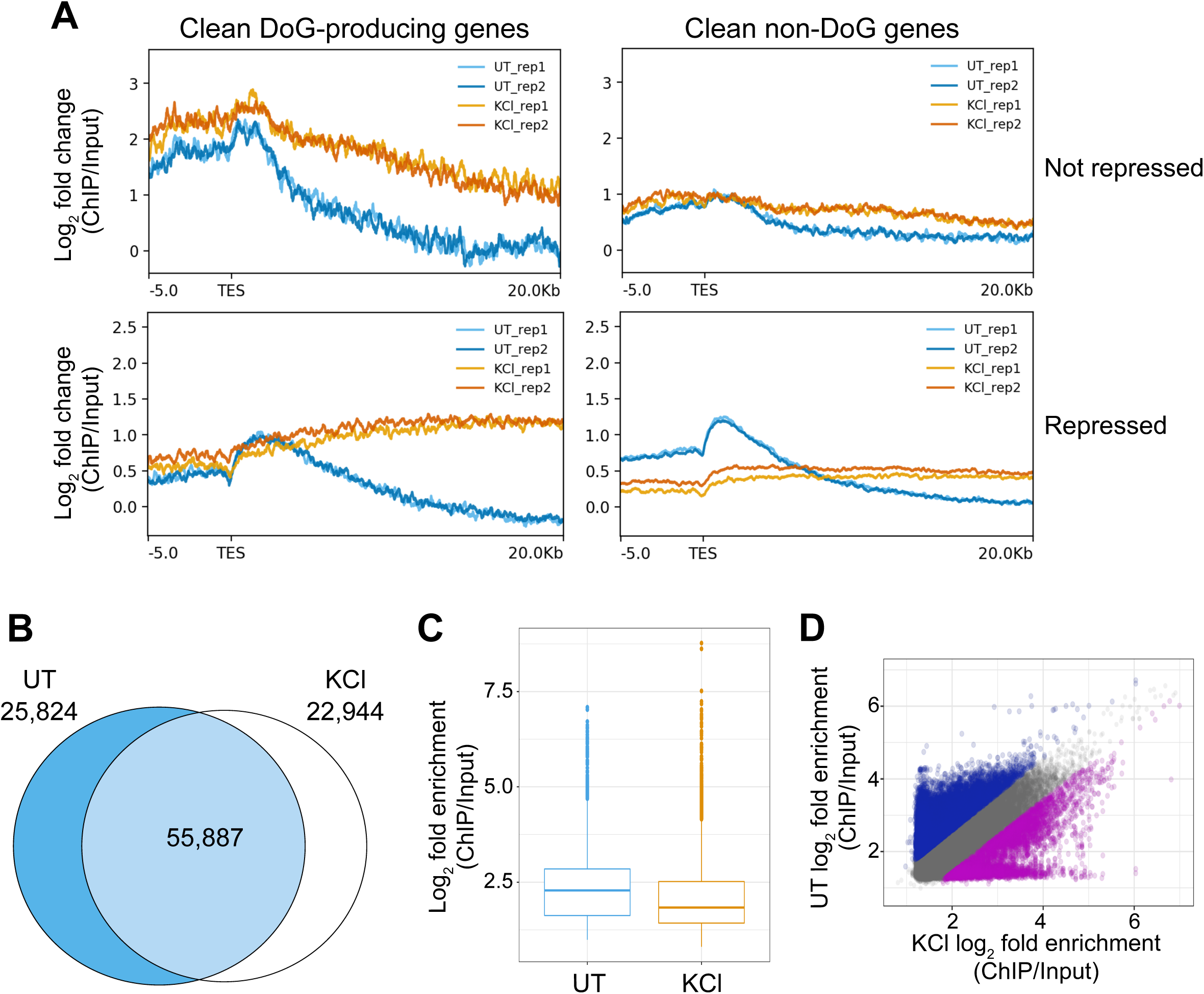
Pol II is redistributed along the genome upon hyperosmotic stress. A) Meta-plots showing the log_2_ fold change of anti-Pol II ChIP-seq data normalized to input across the annotated transcription end sites (TES) for DoG-producing clean genes and clean non-DoG genes that are not repressed (top) and for clean genes that are repressed upon hyperosmotic stress (bottom). Two biological replicates from untreated and KCl-treated HEK-293T cells are shown in each plot. B) Venn diagram showing Pol II binding peaks detected in UT and KCl samples (55,887). 22,944 peaks were detected exclusively in the KCl samples, while 25,824 peaks were found only in the UT samples. C) Box plot depicting the log_2_ fold enrichment (ChIP/Input) of Pol II peaks in UT and KCl treated samples. D) Scatter plot showing the fold enrichment (ChIP/Input) for Pol II binding peaks detected in untreated (y-axis) and KCl-treated samples (x-axis). Data corresponding to peaks that show a decrease in Pol II binding after stress (log2 FC < −0.58) are shown in blue and data for peaks showing increased Pol II binding after stress are shown in purple.

**Figure S5:**
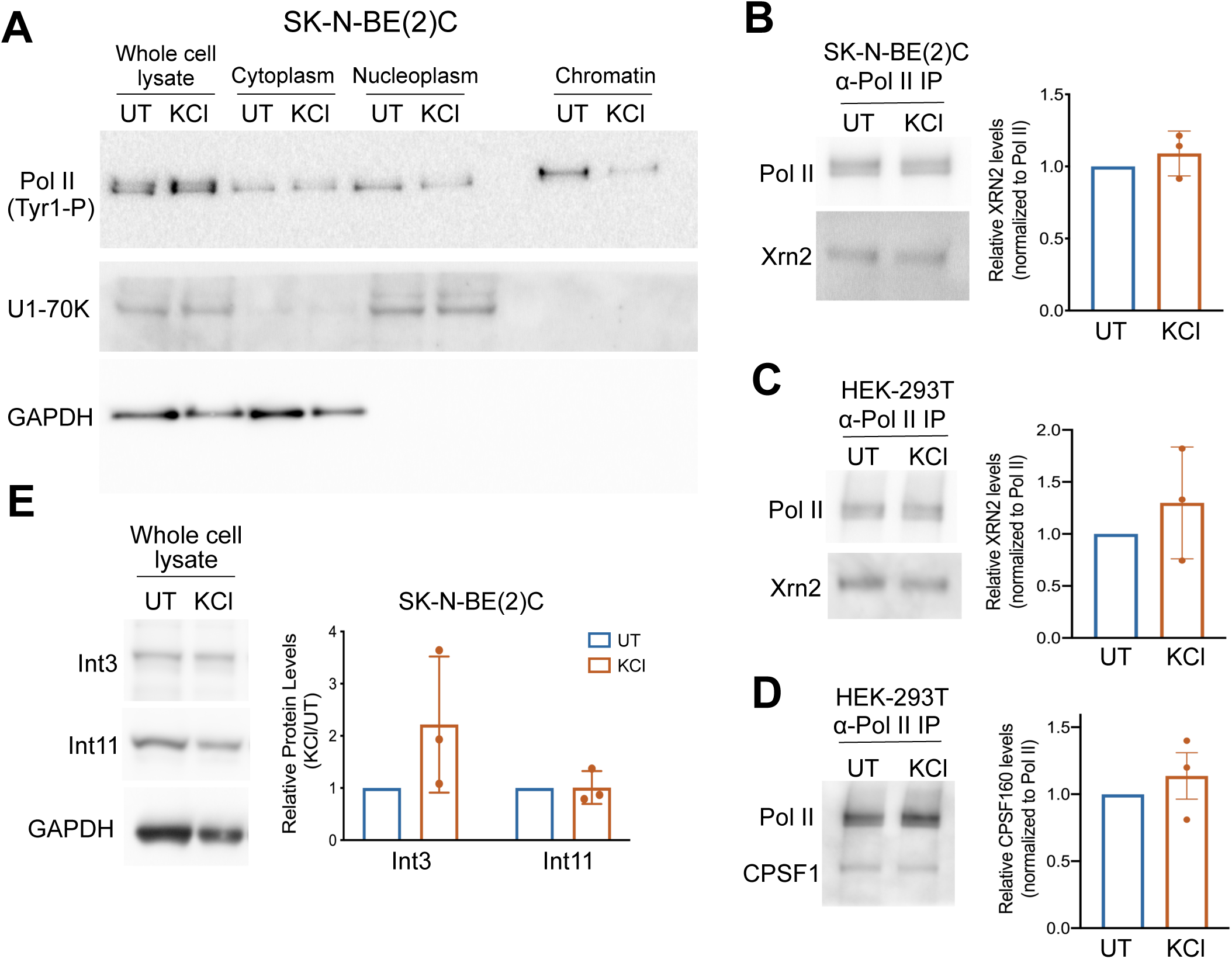
Levels of termination factors remain unchanged after hyperosmotic stress. A) Western blot showing the efficiencies of steps in chromatin fractionation for SK-N-BE(2)C cells. Pol II phosphorylated on Tyr1 residues of the Pol II CTD served as a chromatin marker, U1-70K as a nucleoplasmic marker and GAPDH as a cytoplasmic marker. B, C) Western blot of anti-Pol II immunoprecipitates from B) SK-N-BE(2)C cells and C) HEK-293T cells confirm that Pol II binding to Xrn2 does not decrease after hyperosmotic stress. D) Western blotting of anti-Pol II immunoprecipitates using an antibody against CPSF1 confirms that hyperosmotic stress does not alter the binding between CPSF1 and Pol II. E) Western blot of SK-N-BE(2)C whole cell lysates detect comparable levels of Int3 and Int11 in untreated and KCl-treated samples.

**Figure S6:**
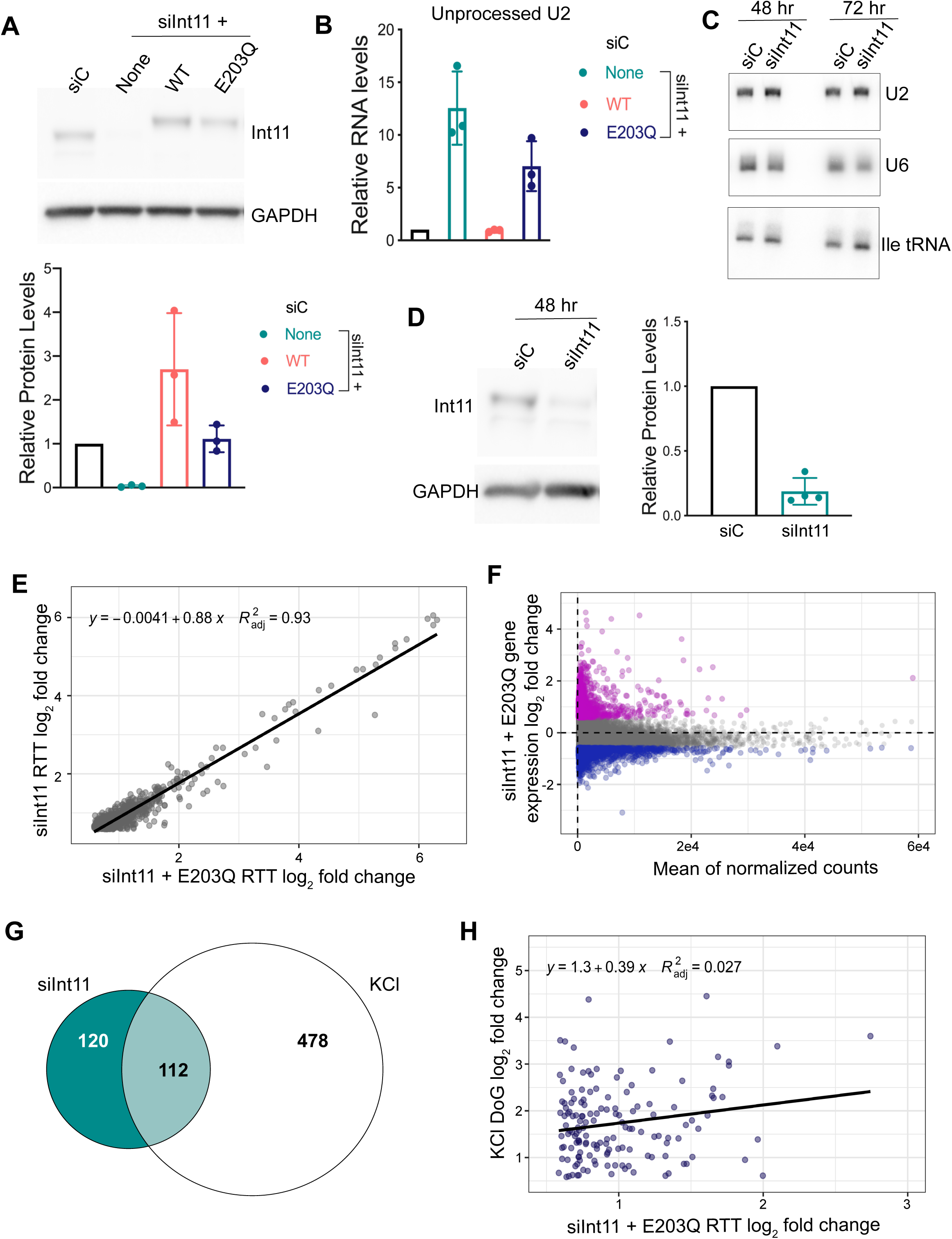
Int11 knockdown leads to readthrough transcription. A) Western blots showing the levels of Int11 and GAPDH in lysates from HEK-293T cells transfected with a scrambled siRNA control (siC) or with an siRNA against Int11 and in rescue cell lines stably expressing WT or E203Q Int11 transfected with an siRNA against Int11 for 72 hours. Bar graph shows the quantification of Int11 band intensities normalized to GAPDH band intensities. B) RT-qPCR results confirm accumulation of unprocessed U2 in siInt11-transfected cells with no rescue (None) or with E203Q rescue, but not in cells with wild-type rescue (WT). C) Probes against U2 snRNA, U6 snRNA and Isoleucine tRNA were used for northern blotting of total RNAs extracted from siC and siInt11-transfected samples 48 hours and 72 hours after transfection. D) Western blots showing the levels of Int11 and GAPDH in lysates from HEK-293T cells transfected with a scrambled siRNA control (siC) or siRNA against Int11 (siInt11) for 48 hours (81% knockdown efficiency). E) Scatter plot showing correlation between readthrough transcripts detected in siInt11-transcfected cells expressing no rescue and cells expressing the E203Q mutant Int11 (n=576). F) Mean average plot showing the average read counts for expressed genes on the x-axis and their log_2_ FC after depletion of functional Int11 (E203Q sample) on the y-axis (n=14,640). Activated genes are shown in purple and repressed genes in blue. G) Venn diagram showing the overlap between the identities of clean genes that produce readthrough transcripts in siInt11-transfected cells (teal) and DoG-producing genes in KCl-treated samples (white). H) Scatter plot demonstrating the correlation between the log_2_ FC of overlapping genes in KCl-treated cells (y-axis) and siInt11 + E203Q cells (x-axis) (n=148).

<u>Table S1</u>: Differential expression analysis of clean genes and DoG-producing clean genes.

<u>Table S2</u>: Gene ontology analysis for DoG-producing clean genes and clean non-DoG genes.

<u>Table S3</u>: Proteins identified by mass spectrometry in untreated and KCl-treated samples.

<u>Table S4</u>: Readthrough transcripts induced upon depletion of functional Int11.

**Table S5:**
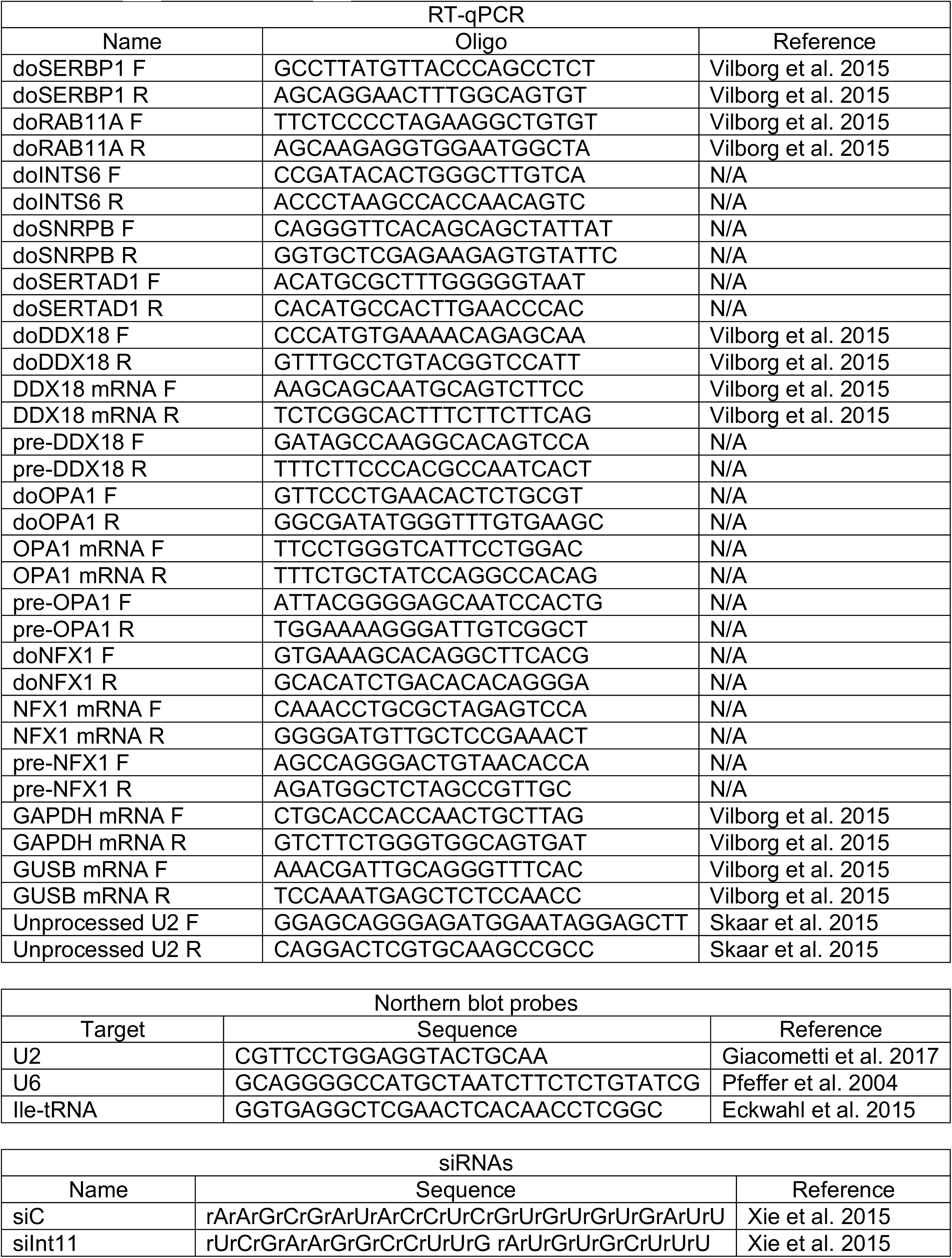
List of oligos used in this study.

## Methods

### Cell culture and inductions

HEK-293T cells were cultured in DMEM supplemented with 10% fetal bovine serum, 1% penicillin and streptomycin, and 1% L-glutamine. Hyperosmotic stress inductions were performed using 80 mM KCl for 1 hour, as described in Vilborg et al. 2015. Stable cell lines expressing WT or E203Q mutant Int11 were cultured as previously described (Xie et al. 2015).

### RNA preparations and RT-qPCRs

RNA extractions were performed using TRIzol according to the manufacturer’s instructions. After treating samples with RQ1 DNase (Promega), RNA was recovered through PCA extraction followed by ethanol precipitation. cDNA was made from 2 µg of RNA using Super Script III and random primers (Invitrogen). Samples were then diluted 1:10 and 2 µL of diluted cDNA were used for RT-qPCR along with 5 µL of iTaq universal SYBR mix (Bio-Rad), 1.5 µL of primers (1.5 µM mix of forward and reverse) and 1.5 µL of water. Plates were run on a Bio-Rad CFX384 machine. Results were analyzed using the comparative C_T_ method (Schmittgen and Livak. 2008). Primers designed for this study were subjected to a primer efficiency test and amplicons were run on agarose gels to ensure that a single band was observed for each pair. All primers used in this study are listed in Table S5.

### High throughput sequencing

#### TT-TimeLapse

Samples were fed 1 mM s^4^U (Sigma) during the final 5 minutes of incubation with either KCl or siRNAs. After extracting RNA from HEK-293T cells, samples were spiked-in with 4% s^4^U-labeled RNA from *Drosophila* S2 cells. TT-TL-seq was performed as described in Schofield et al. 2018. Sequencing was done by the Yale Center for Genomic Analysis (YCGA).

#### ChIP-seq

2.5×10^7^ HEK-293T cells were plated in 15 cm dishes (5×10^6^ per dish) and incubated at 37°C overnight. Cells were crosslinked during the last 10 minutes of stress inductions using 1% paraformaldehyde after which they were washed twice with 1x PBS. Chromatin immunoprecipitations were performed as described in Bieberstein et al. 2014 using 30 µL of Pol II antibody (Cell Signaling D8L4Y) and 180 µL of Protein A beads. The YCGA prepared libraries and sequenced the samples on the NovaSeq platform.

#### Bioinformatics

TT-TL-seq: Spiked-in RNA samples from HEK-293T cells were mapped to a combined hg38 and dm6 genome using hisat2 version 2.1.0 (Kim et al. 2015). Bam files for mapped reads were created, sorted and indexed using samtools version 1.9 (Li et al. 2009). Gene counts were generated using bedtools multicov version 2.26.0 (Quinlan et al. 2010). Normalization factors for each sample were calculated in edgeR (Robinson et al. 2010) using read counts mapped to the dm6 genome. Read-in values used to define clean genes were generated using ARTDeco (Roth et al. 2020). DoGs were identified using DoGFinder (Wiesel et al. 2018). Differential expression analysis of normalized read counts in gene bodies and in DoG regions was done using DESeq2 (Love et al. 2014). The list of clean genes is comprised by genes that were expressed by more than 100 read counts after normalization to spike-in, had a read in value ≤ −1 and did not overlap with DoGs from neighboring genes on either strand according to bedtools intersect analysis (Quinlan et al. 2010). EnrichR was used for gene ontology analysis of biological enrichments (Chen et al. 2013; Kuleshov et al. 2016). Visualization of the generated data was achieved using ggplot2 (Wickham 2016) and graphPad prism. Eulerr was used to generate proportional Venn diagrams. Normalized tracks were visualized using IGV.

#### ChIP-seq

ChIP-seq samples were mapped to the hg38 human genome using bowtie2 (Langmead et al. 2009). Bam files and gene counts were generated as described above. Counts were normalized to input using edgeR normalizeChIPtoInput function (Robinson et al. 2010). Normalized bigwig files and meta plots were created using deepTools version 3.3.0 (Ramírez et al. 2016). Normalized tracks were visualized using IGV. MACS2 version 2.1.1 (Zhang et al. 2008) was used to call Pol II binding peaks and determine their fold enrichment over input in untreated and KCl-treated samples.

### Cell fractionation and immunoprecipitations

For cellular fractionation, 4.8×10^6^ cells were plated in three 15 cm dishes and incubated overnight at 37°C. After KCl treatment, cells were washed twice with 1x PBS, scraped on ice and pelleted by centrifugation (1400rpm at 4°C for 5 minutes). Pellets were resuspended using hypotonic lysis buffer for 5 minutes (Nojima et al. 2016). Nuclei were collected by centrifugation (1400rpm at 4°C for 5 minutes) and the cytoplasmic fraction (supernatant) was discarded. Nuclei were incubated in NUN1 and NUN2 (Wuarin and Schibler 1994) on ice for 15 minutes with intermittent vortexing. Samples were then spun down to pellet the chromatin fraction. Chromatin fragments were generated by incubating the chromatin pellets with 2 µL of micrococcal nuclease for 5 minutes at 37°C, 1400 rpm in a thermomixer (MNase-digested samples) or by incubating the pellets with 25U of benzonase for 45 minutes (Benzonase-digested samples). After digestion, the samples were spun down (16,000 g at 4°C for 5 minutes) and the supernatant was transferred into a new 1.5 mL tube.

For immunoprecipitation, 10 µg of anti-Pol II antibody (MABI0601) or 3 µg of anti-Int3 (Bethyl A302-050A) antibody were incubated with 25 µL of magnetic anti-mouse beads (NEB) overnight. The following day, the antibody-conjugated beads were washed twice with 1 mL of NET-2 buffer and resuspended in 25 µL of NET-2 buffer. Chromatin fragments were pre-cleared by incubating them with the IgG-conjugated beads for 3 hours in a rotator at 4°C. The beads were then collected using a magnetic rack and the supernatant was transferred to a 1.5 mL tube containing the anti-Pol II conjugated beads. The samples were incubated for an additional 2 hours in the rotator at 4°C. After collecting the beads using a magnetic rack, the supernatant was discarded and the beads were washed four times using 1 mL of NET-2 buffer. The final wash was done using 1x PBS and the samples were resuspended using 8 µL of 10 mM Tris-HCL pH 8.5.

### Proteomics

#### Digestion

The anti-Pol II immunoprecipitates were solubilized with a 2.5% solution of ALS-110 (Protea) in 50 mM Tris-HCL pH=8.5 with 5 mM EDTA, and 50 mM DTT directly on the beads. Samples were incubated at 95°C for 10 minutes to reduce cystines. Reactions were cooled down on ice for 1 minute, 2 μL of 1 M Tris pH 8.5 were added and cysteines were alkylated by adding 4.7 μL of 100 mM iodoacetamide for 30 minutes in the dark. The reactions were then quenched with the addition of 0.7 μL of 200 mM DTT. Afterwards, 108 μL water, 0.67 μL CaCl_2_, 1.33 μL 1 M Tris-HCL pH 8.5 and 7.8 μL 0.5 mg/mL trypsin (Promega) were added and proteins left to digest for 16 hours at 37°C. Surfactant was cleaved with the addition of 20% trifluoroacetic acid until pH < 3 and samples were incubated at 25°C for 15 minutes. Peptides were then separated from beads using a magnetic rack and desalted using C18 silica MicroSpin Columns (The Nest Group). Column elution was performed in 300 μL 80% acetonitrile 0.1% trifluoroacetic acid and peptides were dried by centrifugal vacuum.

#### Mass spectrometry

LC-MS/MS was performed using an ACQUITY UPLC M-Class (Waters) paired with a Q Exactive Plus (Thermo). Peptides were separated on a 65-cm-long, 75-μm-internal-diameter PicoFrit column (New Objective) packed in-house to a length of 50 cm with 1.9 μm ReproSil-Pur 120 Å C18-AQ (Dr. Maisch) using methanol as the packing solvent. Peptide separation was achieved using a non-linear 90-min gradient from 1% ACN 0.1% formic acid to 90% ACN 0.1% formic acid with a flowrate of 250 nl/min. Approximately 3-4 μg of peptides were run for each of the seven biological replicates with at least one blank run between samples to avoid peptide carryover.

#### Bioinformatics

Mass spectra were searched using Maxquant version 1.5.1.2 (Cox et al. 2008) or MASCOT version 2.6.0.0 (Perkins et al. 1999) with Carbamidomethyl (C) as a fixed modification, Acetyl (N-Term), Deamidation (NQ), Oxidation (M) and Phosphorylation (S/T/Y) as variable modifications and with up to three missed trypsin cleavages, a 5 amino acid minimum length, and 1% false discovery rate (FDR) against the Uniprot Human database (downloaded Dec-2017). Searches were then analyzed with Perseus version 1.5.8.5. The analysis included contaminant and reverse protein hits removal, log_2_ transformation, and averaging across the seven biological replicates.

### siRNA knockdowns

For knockdown experiments, 7.5×10^4^ HEK-293T cells were seeded in six well plates and incubated at 37°C overnight. After 24 hours, the cells were transfected with either 50 nM of either scrambled siRNA control (siC) or siRNA against Int11 (siInt11) (Xie et al. 2015) using Lipofectamine RNAimax according to manufacturer’s instructions. Cells were incubated at 37°C for 72 hours after which they were washed with 1x PBS, scraped and pelleted by centrifugation (1400 rpm at 4°C for 5 minutes). After discarding the supernatant, cells were resuspended in 150 µL of NET-2 buffer. 50 µL were aliquoted to verify knockdown efficiency through western blots and the rest of the sample was lysed using TRIzol. siRNA experiments for TT-TL experiments were performed by transfecting 1.5×10^5^ cells for 48 hours instead of 72 hours.

### Western blots

For western blots of whole cell lysates, samples were sonicated using 10 cycles of 30 seconds on and 30 seconds off in a Diagenode Bioruptor Pico sonicator (Withers et al. 2018) and digested with micrococcal nuclease for 45 minutes at 37°C. Samples were run at an increasing voltage (up to 120 V) in NuPAGE 4-12% BisTris gels in 1x MOPS buffer (Invitrogen). Proteins were transferred overnight to a 0.45 µm nitrocellulose membrane at 30 V. Membranes were blocked for 1 hour in 5% milk in 1x PBST. Detection of total Pol II and Tyr1-P containing Pol II molecules was achieved using antibodies from MBL (MABI0601, discontinued) and Active motif (61383) respectively, at a 1:1000 dilution. Antibodies against Ints3 (Bethyl A302-050A), Ints11 (Abcam ab75276), Xrn2 (A301-101) and CPSF160 (A301-580A) were used at a 1:800 dilution. Anti-GAPDH antibody (Cell Signaling 14C10) was used at 1:2000, while the antibody against U1-70K (Kastner et al. 1992; Tarn and Steitz. 1994) was used at a 1:500 dilution. All secondary antibodies were used at a 1:2000 dilution of 1x PBST, 3% milk.

## References

Albrecht, T.R., and Wagner, E.J. (2012). snRNA 3’ end formation requires heterodimeric association of integrator subunits. Mol Cell Biol 32, 1112–1123.

Amat, R., Bottcher, R., Le Dily, F., Vidal, E., Quilez, J., Cuartero, Y., Beato, M., de Nadal, E., and Posas, F. (2019). Rapid reversible changes in compartments and local chromatin organization revealed by hyperosmotic shock. Genome Res 29, 18–28.

Baillat, D., Hakimi, M.A., Naar, A.M., Shilatifard, A., Cooch, N., and Shiekhattar, R. (2005). Integrator, a multiprotein mediator of small nuclear RNA processing, associates with the C-terminal repeat of RNA Polymerase II. Cell 123, 265–276.

Baillat, D., and Wagner, E.J. (2015). Integrator: surprisingly diverse functions in gene expression. Trends Biochem Sci 40, 257–264.

Bauer, D.L.V., Tellier, M., Martinez-Alonso, M., Nojima, T., Proudfoot, N.J., Murphy, S., and Fodor, E. (2018). Influenza virus mounts a two-pronged attack on host RNA Polymerase II transcription. Cell Rep 23, 2119–2129 e2113.

Bieberstein, N.I., Straube, K., and Neugebauer, K.M. (2014). Chromatin immunoprecipitation approaches to determine co-transcriptional nature of splicing. Methods Mol Biol 1126, 315–323.

Cardiello, J.F., Goodrich, J.A., and Kugel, J.F. (2018). Heat shock causes a reversible increase in RNA Polymerase II occupancy downstream of mRNA genes, consistent with a global loss in transcriptional termination. Mol Cell Biol 38.

Chen, E.Y., Tan, C.M., Kou, Y., Duan, Q., Wang, Z., Meirelles, G.V., Clark, N.R., and Ma’ayan, A. (2013). Enrichr: interactive and collaborative HTML5 gene list enrichment analysis tool. BMC Bioinformatics 14, 128

Chen, K., Hu, Z., Xia, Z., Zhao, D., Li, W., and Tyler, J.K. (2015). The overlooked fact: fundamental need for spike-in control for virtually all genome-wide analyses. Mol Cell Biol 36, 662–667.

Cox, J., & Mann, M. (2008). MaxQuant enables high peptide identification rates, individualized p.p.b.-range mass accuracies and proteome-wide protein quantification. Nat. Biotechnol. 26, 1367–1372.

Dominski, Z., Yang, X.C., Purdy, M., Wagner, E.J., and Marzluff, W.F. (2005). A CPSF-73 homologue is required for cell cycle progression but not cell growth and interacts with a protein having features of CPSF-100. Mol Cell Biol 25, 1489–1500.

Duffy, E.E., Rutenberg-Schoenberg, M., Stark, C.D., Kitchen, R.R., Gerstein, M.B., and Simon, M.D. (2015). Tracking distinct RNA populations using efficient and reversible covalent chemistry. Mol Cell 59, 858–866.

Eaton, J.D., Davidson, L., Bauer, D.L.V., Natsume, T., Kanemaki, M.T., and West, S. (2018). Xrn2 accelerates termination by RNA Polymerase II, which is underpinned by CPSF73 activity. Genes Dev 32, 127–139.

Eaton, J.D., Francis, L., Davidson, L., and West, S. (2020). A unified allosteric/torpedo mechanism for transcriptional termination on human protein-coding genes. Genes Dev 34, 132–145.

Elrod, N.D., Henriques, T., Huang, K.L., Tatomer, D.C., Wilusz, J.E., Wagner, E.J., and Adelman, K. (2019). The Integrator complex attenuates promoter-proximal transcription at protein-coding genes. Mol Cell 76, 738–752.e737.

Fong, N., Brannan, K., Erickson, B., Kim, H., Cortazar, M.A., Sheridan, R.M., Nguyen, T., Karp, S., and Bentley, D.L. (2015). Effects of transcription elongation rate and Xrn2 exonuclease activity on RNA Polymerase II termination suggest widespread kinetic competition. Mol Cell 60, 256–267.

Gardini, A., Baillat, D., Cesaroni, M., Hu, D., Marinis, J.M., Wagner, E.J., Lazar, M.A., Shilatifard, A., and Shiekhattar, R. (2014). Integrator regulates transcriptional initiation and pause release following activation. Mol Cell 56, 128–139.

Grosso, A.R., Leite, A.P., Carvalho, S., Matos, M.R., Martins, F.B., Vitor, A.C., Desterro, J.M., Carmo-Fonseca, M., and de Almeida, S.F. (2015). Pervasive transcription read-through promotes aberrant expression of oncogenes and RNA chimeras in renal carcinoma. Elife 4.

Harlen, K.M., Trotta, K.L., Smith, E.E., Mosaheb, M.M., Fuchs, S.M., and Churchman, L.S. (2016). Comprehensive RNA Polymerase II interactomes reveal distinct and varied roles for each phospho-CTD residue. Cell Rep 15, 2147–2158.

Heinz, S., Texari, L., Hayes, M.G.B., Urbanowski, M., Chang, M.W., Givarkes, N., Rialdi, A., White, K.M., Albrecht, R.A., Pache, L., et al. (2018). Transcription elongation can affect genome 3D structure. Cell 174, 1522–1536.e1522.

Hennig, T., Michalski, M., Rutkowski, A.J., Djakovic, L., Whisnant, A.W., Friedl, M.S., Jha, B.A., Baptista, M.A.P., L’Hernault, A., Erhard, F., et al. (2018). HSV-1-induced disruption of transcription termination resembles a cellular stress response but selectively increases chromatin accessibility downstream of genes. PLoS Pathog 14, e1006954.

Jalihal, A.P., Pitchiaya, S., Xiao, L., Bawa, P., Jiang, X., Bedi, K., Parolia, A., Cieslik, M., Ljungman, M., Chinnaiyan, A.M., et al. (2020). Multivalent proteins rapidly and reversibly phase-separate upon osmotic cell volume change. bioRxiv, 748293.

Kastner, B., Kornstadt, U., Bach, M., and Luhrmann, R. (1992). Structure of the small nuclear RNP particle U1: identification of the two structural protuberances with RNP-antigens A and 70K. J Cell Biol 116, 839–849.

Kim, D., Langmead, B., and Salzberg, S.L. (2015). HISAT: a fast spliced aligner with low memory requirements. Nat Methods 12, 357–360.

Kuleshov, M.V., Jones, M.R., Rouillard, A.D., Fernandez, N.F., Duan, Q., Wang, Z., Koplev, S., Jenkins, S.L., Jagodnik, K.M., Lachmann, A., et al. (2016). Enrichr: a comprehensive gene set enrichment analysis web server 2016 update. Nucleic Acids Res 44, W90–97.

Lai, F., Gardini, A., Zhang, A., and Shiekhattar, R. (2015). Integrator mediates the biogenesis of enhancer RNAs. Nature 525, 399–403.

Langmead, B., Trapnell, C., Pop, M., and Salzberg, S.L. (2009). Ultrafast and memory-efficient alignment of short DNA sequences to the human genome. Genome Biol 10, R25.

Li, H., Handsaker, B., Wysoker, A., Fennell, T., Ruan, J., Homer, N., Marth, G., Abecasis, G., Durbin, R., and Genome Project Data Processing, S. (2009). The sequence alignment/map format and SAMtools. Bioinformatics 25, 2078–2079.

Love, M.I., Huber, W., and Anders, S. (2014). Moderated estimation of fold change and dispersion for RNA-seq data with DESeq2. Genome Biol 15, 550.

Lovén, J., Orlando, D.A., Sigova, A.A., Lin, C.Y., Rahl, P.B., Burge, C.B., Levens, D.L., Lee, T.I., and Young, R.A. (2012). Revisiting global gene expression analysis. Cell 151, 476–482.

Mahat, D.B., Salamanca, H.H., Duarte, F.M., Danko, C.G., and Lis, J.T. (2016). Mammalian heat shock response and mechanisms underlying its genome-wide transcriptional regulation. Mol Cell 62, 63–78.

Mak, S.K., and Kültz, D. (2004). Gadd45 proteins induce G2/M arrest and modulate apoptosis in kidney cells exposed to hyperosmotic stress. J Biol Chem 279, 39075–39084.

Mandel, C.R., Kaneko, S., Zhang, H., Gebauer, D., Vethantham, V., Manley, J.L., and Tong, L. (2006). Polyadenylation factor CPSF-73 is the pre-mRNA 3’-end-processing endonuclease. Nature 444, 953–956.

Mazo, A., Hodgson, J.W., Petruk, S., Sedkov, Y., and Brock, H.W. (2007). Transcriptional interference: an unexpected layer of complexity in gene regulation. J Cell Sci 120, 2755–2761.

Muniz, L., Deb, M.K., Aguirrebengoa, M., Lazorthes, S., Trouche, D., and Nicolas, E. (2017). Control of gene expression in senescence through transcriptional read-through of convergent protein-coding genes. Cell Rep 21, 2433–2446.

Nemeroff, M.E., Barabino, S.M., Li, Y., Keller, W., and Krug, R.M. (1998). Influenza virus NS1 protein interacts with the cellular 30 kDa subunit of CPSF and inhibits 3’end formation of cellular pre-mRNAs. Mol Cell 1, 991–1000.

Nojima, T., Gomes, T., Carmo-Fonseca, M., and Proudfoot, N.J. (2016). Mammalian NET-seq analysis defines nascent RNA profiles and associated RNA processing genome-wide. Nat Protoc 11, 413–428.

Nojima, T., Tellier, M., Foxwell, J., Ribeiro de Almeida, C., Tan-Wong, S.M., Dhir, S., Dujardin, G., Dhir, A., Murphy, S., and Proudfoot, N.J. (2018). Deregulated expression of mammalian lncRNA through loss of SPT6 induces R-Loop formation, replication stress, and cellular senescence. Mol Cell 72, 970–984.e977.

Perkins, D.N., Pappin, D.J., Creasy, D.M., Cottrell, J.S. (1999). Probability-based protein identification by searching sequence databases using mass spectrometry data. Electrophoresis. 20, 3551–67.

Proudfoot, N.J. (2016). Transcriptional termination in mammals: Stopping the RNA Polymerase II juggernaut. Science 352, aad9926.

Quinlan, A.R., and Hall, I.M. (2010). BEDTools: a flexible suite of utilities for comparing genomic features. Bioinformatics 26, 841–842.

Ramírez, F., Ryan, D.P., Gruning, B., Bhardwaj, V., Kilpert, F., Richter, A.S., Heyne, S., Dundar, F., and Manke, T. (2016). deepTools2: a next generation web server for deep-sequencing data analysis. Nucleic Acids Res 44, W160–165.

Robbins, E., Pederson, T., and Klein, P. (1970). Comparison of mitotic phenomena and effects induced by hypertonic solutions in HeLa cells. J Cell Biol 44, 400–416.

Robinson, M.D., McCarthy, D.J., and Smyth, G.K. (2010). edgeR: a Bioconductor package for differential expression analysis of digital gene expression data. Bioinformatics 26, 139–140.

Roth, S.J., Heinz, S., and Benner, C. (2020). ARTDeco: automatic readthrough transcription detection. BMC Bioinformatics 21, 214.

Rutkowski, A.J., Erhard, F., L’Hernault, A., Bonfert, T., Schilhabel, M., Crump, C., Rosenstiel, P., Efstathiou, S., Zimmer, R., Friedel, C.C., et al. (2015). Widespread disruption of host transcription termination in HSV-1 infection. Nat Commun 6, 7126.

Schmittgen, T.D., and Livak, K.J. (2008). Analyzing real-time PCR data by the comparative C(T) method. Nat Protoc 3, 1101–1108.

Schofield, J.A., Duffy, E.E., Kiefer, L., Sullivan, M.C., and Simon, M.D. (2018). TimeLapse-seq: adding a temporal dimension to RNA sequencing through nucleoside recoding. Nat Methods 15, 221–225.

Schwalb, B., Michel, M., Zacher, B., Fruhauf, K., Demel, C., Tresch, A., Gagneur, J., and Cramer, P. (2016). TT-seq maps the human transient transcriptome. Science 352, 1225–1228.

Shalgi, R., Hurt, J.A., Lindquist, S., and Burge, C.B. (2014). Widespread inhibition of posttranscriptional splicing shapes the cellular transcriptome following heat shock. Cell Rep 7, 1362–1370.

Shi, Y., Di Giammartino, D.C., Taylor, D., Sarkeshik, A., Rice, W.J., Yates, J.R., 3rd, Frank, J., and Manley, J.L. (2009). Molecular architecture of the human pre-mRNA 3’ processing complex. Mol Cell 33, 365–376.

Skaar, J.R., Ferris, A.L., Wu, X., Saraf, A., Khanna, K.K., Florens, L., Washburn, M.P., Hughes, S.H., and Pagano, M. (2015). The Integrator complex controls the termination of transcription at diverse classes of gene targets. Cell Res 25, 288–305.

Stadelmayer, B., Micas, G., Gamot, A., Martin, P., Malirat, N., Koval, S., Raffel, R., Sobhian, B., Severac, D., Rialle, S., et al. (2014). Integrator complex regulates NELF-mediated RNA Polymerase II pause/release and processivity at coding genes. Nat Commun 5, 5531.

Tarn, W.Y., and Steitz, J.A. (1994). SR proteins can compensate for the loss of U1 snRNP functions in vitro. Genes Dev 8, 2704–2717.

Tatomer, D.C., Elrod, N.D., Liang, D., Xiao, M.S., Jiang, J.Z., Jonathan, M., Huang, K.L., Wagner, E.J., Cherry, S., and Wilusz, J.E. (2019). The Integrator complex cleaves nascent mRNAs to attenuate transcription. Genes Dev 33, 1525–1538.

Vilborg, A., Passarelli, M.C., Yario, T.A., Tycowski, K.T., and Steitz, J.A. (2015). Widespread inducible transcription downstream of human genes. Mol Cell 59, 449–461.

Vilborg, A., Sabath, N., Wiesel, Y., Nathans, J., Levy-Adam, F., Yario, T.A., Steitz, J.A., and Shalgi, R. (2017). Comparative analysis reveals genomic features of stress-induced transcriptional readthrough. Proc Natl Acad Sci U S A 114, E8362–E8371.

Vilborg, A., and Steitz, J.A. (2017). Readthrough transcription: How are DoGs made and what do they do? RNA Biol 14, 632–636.

Wang, X., Hennig, T., Whisnant, A.W., Erhard, F., Prusty, B.K., Friedel, C.C., Forouzmand, E., Hu, W., Erber, L., Chen, Y., et al. (2020). Herpes simplex virus blocks host transcription termination via the bimodal activities of ICP27. Nat Commun 11, 293.

Wickham, H. (2016). ggplot2: Elegant Graphics for Data Analysis (New York:Springer-Verlag).

Wiesel, Y., Sabath, N., and Shalgi, R. (2018). DoGFinder: a software for the discovery and quantification of readthrough transcripts from RNA-seq. BMC Genomics 19, 597.

Withers, J.B., Li, E.S., Vallery, T.K., Yario, T.A., and Steitz, J.A. (2018). Two herpesviral noncoding PAN RNAs are functionally homologous but do not associate with common chromatin loci. PLoS Pathog 14, e1007389.

Wu, Y., Albrecht, T.R., Baillat, D., Wagner, E.J., and Tong, L. (2017). Molecular basis for the interaction between Integrator subunits IntS9 and IntS11 and its functional importance. Proc Natl Acad Sci U S A 114, 4394–4399.

Wuarin, J., and Schibler, U. (1994). Physical isolation of nascent RNA chains transcribed by RNA Polymerase II: evidence for cotranscriptional splicing. Mol Cell Biol 14, 7219–7225.

Xie, M., Zhang, W., Shu, M.D., Xu, A., Lenis, D.A., DiMaio, D., and Steitz, J.A. (2015). The host Integrator complex acts in transcription-independent maturation of herpesvirus microRNA 3’ ends. Genes Dev 29, 1552–1564.

Zhang, Y., Liu, T., Meyer, C.A., Eeckhoute, J., Johnson, D.S., Bernstein, B.E., Nusbaum, C., Myers, R.M., Brown, M., Li, W., et al. (2008). Model-based analysis of ChIP-Seq (MACS). Genome Biol 9, R137.

## Additional references

Giacometti, S., Benbahouche, N.E.H., Domanski, M., Robert, M.C., Meola, N., Lubas, M., Bukenborg, J., Andersen, J.S., Schulze, W.M., Verheggen, C., et al. (2017). Mutually Exclusive CBC-Containing Complexes Contribute to RNA Fate. Cell Rep 18, 2635–2650.

Pfeffer, S., Zavolan, M., Grässer, F.A., Chien, M., Russo, J.J., Ju, J., John, B., Enright, A.J., Marks, D., Sander, C., et al. (2004). Identification of virus-encoded microRNAs. Science 304, 734–736.

Eckwahl, M.J., Sim, S., Smith, D., Telesnitsky, A., and Wolin, S.L. (2015). A retrovirus packages nascent host noncoding RNAs from a novel surveillance pathway. Genes Dev 29, 646–657.

